# Selective targeting of TBXT with DARPins identifies regulatory networks and therapeutic vulnerabilities in chordoma

**DOI:** 10.1101/2024.09.20.614025

**Authors:** Charles S. Umbaugh, Marie Groth, Cihan Erkut, Kwang-Seok Lee, Joana Marinho, Florian Iser, Jonas N. Kapp, Petra Schroeter, Simay Dolaner, Asli Kayserili, Julia Hartmann, Philipp Walch, Thomas F.E. Barth, Kevin Mellert, Birgit Dreier, Jonas V. Schaefer, Andreas Plückthun, Stefan Fröhling, Claudia Scholl

**Affiliations:** Division of Applied Functional Genomics, German Cancer Research Center (DKFZ), Heidelberg, Germany; National Center for Tumor Diseases (NCT), NCT Heidelberg, a partnership between DKFZ and Heidelberg University Hospital, Heidelberg, Germany; Division of Translational Medical Oncology, DKFZ, Heidelberg, Germany; Faculty of Biosciences, Heidelberg University, Heidelberg, Germany; German Cancer Consortium (DKTK), Heidelberg, Germany; Department of Biochemistry, University of Zurich, Zurich, Switzerland; Institute of Pathology, Ulm University, Ulm, Germany; Institute of Human Genetics, Heidelberg University, Heidelberg, Germany

## Abstract

Aberrant expression of the embryonal transcription factor TBXT (also known as brachyury) drives chordoma, a rare spinal neoplasm with no effective drug therapies. The gene network regulated by TBXT is poorly understood, and strategies to disrupt its abnormal activity for therapeutic purposes are lacking. Here, we developed TBXT-targeted designed ankyrin repeat proteins (T-DARPins) that selectively bind TBXT, inhibiting its binding to DNA and expression. In chordoma cells, T-DARPins reduced cell cycle progression, spheroid formation, and tumor growth in mice and induced morphologic changes indicative of senescence and differentiation. Combining T-DARPin-mediated TBXT inhibition with transcriptomic and proteomic analyses, we determined the TBXT regulome in chordoma cells, which comprises in particular networks involved in cell cycle regulation, DNA replication and repair, embryonal cell identity, metabolic processes, and interferon response. The analysis of selected TBXT regulome components provided new insights into chordoma biology, such as the strong upregulation of IGFBP3 upon TBXT inhibition to fine-tune part of TBXT’s downstream effectors. Finally, we assigned each TBXT regulome member a druggability status to create a resource for future translational studies and found high interferon response signaling in chordoma cell lines and patient tumors, which was promoted by TBXT and associated with strong sensitivity to clinically approved JAK2 inhibitors. These findings demonstrate the potential of DARPins to investigate the function of nuclear proteins to understand the regulatory networks of cancers driven by aberrant transcription factor activity, including novel entry points for targeted therapies that warrant testing in patients.

## INTRODUCTION

Chordoma is a rare cancer that typically arises at the sacrum, mobile spine, or skull base, likely from remnants of the notochord, and accounts for up to 4% of primary bone tumors and 20% of primary spine tumors [1]. Clinical characteristics include locally aggressive growth, a tendency to recur, the potential for metastasis, and resistance to conventional chemotherapy [2]. First-line treatment is usually surgery with the goal of complete resection, followed by adjuvant radiotherapy. However, local control is rarely achieved since most chordomas are adjacent to vital structures, particularly at the skull base. As local therapies are exhausted in most patients as the disease progresses, novel approaches to systemic treatment of advanced chordoma are urgently needed.

A potential therapeutic target is the *TBXT* gene (also known as brachyury), which is amplified in 7%, duplicated in 27%, and overexpressed in more than 90% of cases [3, 4] and has been identified as the key driver of chordoma development and progression [3, 5, 6]. It encodes a T-box family transcription factor that regulates mesoderm and notochord formation and axial skeleton development during embryogenesis but is not expressed in most adult tissues [7, 8]. TBXT exerts transcriptional activity by binding as a homodimer to T-box element-containing DNA in the promoter or enhancer regions of target genes [9, 10]. TBXT itself is expressed under the control of a super-enhancer, regulates its own expression, and can control other genes such as *SOX9*, *SAE1*, *TPX2*, and *ATP6V1B2* through direct binding to other super-enhancers [5].

Previous efforts to identify targeted chordoma drug treatments have, for example, focused on mTOR, PI3K, CDK4/6, and EGFR inhibition [11–13]. However, the clinical efficacy of these approaches varies[14, 15]. To suppress *TBXT* transcription, several strategies have been explored preclinically, e.g., inhibition of the enhancer-associated components CDK7/12/13 [5, 6], the receptor tyrosine kinase EGFR [16], or H3K27-specific demethylases [17]. Other approaches to indirect TBXT targeting include vaccines [18, 19] and degradation via bifunctional TBXT-binding nucleic acids that recruit an E3 ligase [20]. Direct pharmacologic inhibition of TBXT is complicated by the fundamental architecture of transcription factors, which impedes direct binding of small molecules to functionally important domains [21]. However, a novel derivative of the EGFR inhibitor afatinib, DHC-156, provides the first proof of principle for direct TBXT inhibition [22].

A novel class of molecules with the potential to inhibit “undruggable” proteins are designed ankyrin repeat proteins (DARPins), consisting of a variable number of internal repeats with a randomized surface that mediates tight and specific binding to the target protein, flanked by capping repeats with a hydrophilic surface [23]. They functionally resemble monoclonal antibodies at only about 10% of the molecular mass, are readily produced in routine bacterial cultures, and can be fused to generate multivalent proteins that bind multiple epitopes or adjacent target proteins [24–26]. DARPins have demonstrated their potential as specific and effective therapeutics targeting extracellular proteins. For example, abicipar pegol inhibits the function of VEGF as effectively as an established Fab fragment of a VEGF-directed monoclonal antibody but with potentially longer duration [27, 28]. In contrast to antibodies, DARPins fold in the cytoplasm and can thus be developed for intracellular applications, where tightly regulated expression is achieved by delivering DARPin-encoding DNA via viral vectors with inducible and/or tissue-specific promoters [29] [30]. A recent example is the inhibition of KRAS isoforms that drive oncogenic signaling in the cytoplasm of cancer cells [31]. However, it has not yet been investigated whether DARPins can also block the function of nuclear proteins, whose deregulation underlies many hematologic and solid-organ malignancies.

In this study, we developed and preclinically characterized DARPins that bind TBXT, thereby blocking its interaction with DNA. Furthermore, we tested these compounds in chordoma cells to determine their utility for direct targeting of an essential transcription factor, explore the phenotypic and transcriptional consequences of TBXT inhibition, and identify components of the gene network regulated by TBXT that could be new entry points for therapeutic intervention.

## RESULTS

### Development of DARPins that block the TBXT-DNA interaction

To generate DARPins that prevent TBXT from binding to DNA, we performed ribosome display selection with a library of ∼10^12^ DARPins against the DNA binding domain (DBD) of TBXT, which yielded 380 candidates with varying specificity and affinity (**Fig. 1a**). These were further narrowed down to 23 potential TBXT-targeting DARPins based on monomeric behavior in size-exclusion chromatography, signal intensities in enzyme-linked immunosorbent assays (ELISA), and homogenous time-resolved fluorescence (HTRF) using biotinylated DBD and full-length TBXT proteins (**Fig. 1a–c; Supplementary Fig. 1a and b**). The selection criteria were binding to both the DBD and full-length TBXT and high but varying HTRF signals, which indicate different binding geometries and thus increase the chances of obtaining different binding modes.

**Figure 1.**
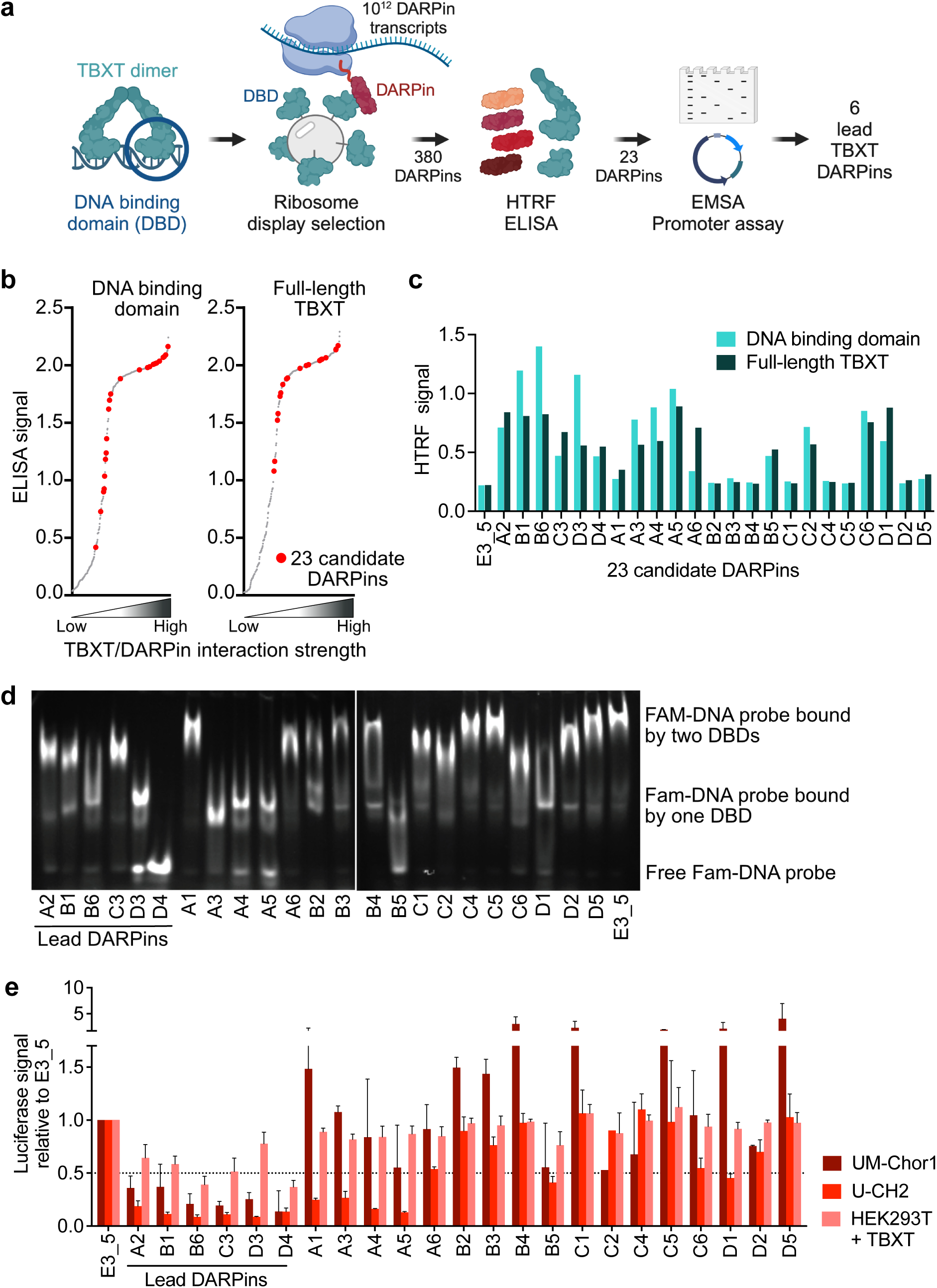
Generation of T-DARPins. **(a)** Schematic of the screening, selection, and testing of DARPins binding the TBXT DBD. Created in BioRender. Fröhling, S. (2024) BioRender.com/l53i374. **(b)** ELISA signal intensities (y-axis) with purified full-length TBXT (right) and the DBD (left) against crude bacterial extracts of the 380 DARPins retrieved by ribosome display selection, ordered by signal strength (x-axis). The 23 candidate DARPins are shown in red. **(c)** HTRF signal for binding of the non-targeting control DARPin E3_5 and the 23 candidate DARPins to the TBXT DBD and full-length TBXT. **(d)** EMSA with a 6-FAM-labeled DNA probe containing two palindromic TBXT binding motifs, recombinant TBXT DBD, and the 23 candidate DARPins or E3_5. **(e)** Dual-luciferase assay with a TBXT-responsive reporter vector in UM-Chor1, U-CH2, and HEK293T cells stably expressing HA-tagged TBXT and lentivirally transduced with 23 candidate TBXT DARPins or E3_5. The dotted line indicates a 50% luciferase signal relative to control. Mean + SEM of two to five biological replicates: UM-Chor1 and U-CH2 with E3_5, A2, B1, B6, C3, or D4, n = 5; UM-Chor1 and U-CH2 with the remaining DARPins, n = 2; TBXT-expressing HEK293 with C2 and D5, n = 2; TBXT-expressing HEK293 with the remaining DARPins, n = 3.

To directly measure the capability of the 23 DARPins to disrupt TBXT binding to DNA, we performed electrophoretic mobility shift assays (EMSA) using a 6-carboxyfluorescein (6-FAM)- labeled DNA probe containing two palindromic TBXT binding sites and recombinant DBD protein. DNA bound by two TBXT DBDs is less mobile during gel electrophoresis and thus shifted to the top. In contrast, unbound DNA or DNA partially occupied by a TBXT DBD moves faster through the gel and is detected in the middle or at the bottom, respectively (**Fig. 1d**). We observed DARPins that largely failed to disrupt DBD-DNA complexes (e.g., A1, C4, C5, and D5) and behaved like the non-targeting control DARPin E3_5 [32]. We also found DARPins with a moderate ability to disrupt the interaction between the DBD and one or two TBXT binding sites (e.g., A2, B1, B6, C3, D3, A3, A4, A5, and D1), and one DARPin (D4) that caused complete displacement of the DNA probe (**Fig. 1d**).

Finally, we measured the candidate DARPins’ capacity to inhibit TBXT transcriptional activity. For this, we generated a reporter vector containing two TBXT response elements upstream of a minimal promoter that drives the expression of *Firefly* luciferase upon binding of TBXT. Using dual-luciferase reporter assays in U-CH2 and UM-Chor1 chordoma cells expressing endogenous TBXT, six of 23 candidate DARPins blocked TBXT binding to the response elements, as evidenced by a reduction in the luciferase signal by more than 50% relative to E3_5 (**Fig. 1e**). These lead DARPins also reduced TBXT binding to the response elements in TBXT-negative HEK293 cells expressing exogenous HA-tagged TBXT (**Fig. 1e**). Thus, we identified six DARPins, referred to as T-DARPins, that bind TBXT at its DBD, thereby displacing it from DNA and preventing it from binding to TBXT-specific transcription factor binding motifs in cells.

### T-DARPins are specific for TBXT and bind to disease-related TBXT variants

Next, we asked whether the T-DARPins effectively and specifically bind endogenous TBXT in chordoma cells. Immunoprecipitation (IP) of Flag-tagged T-DARPins with an anti-Flag antibody from U-CH2 cell lysate confirmed the binding of TBXT to T-DARPins A2, B1, B6, C3, D3, and D4, whereas E3_5 did not pull down TBXT (**Fig. 2a**). In the corresponding whole-cell lysates, we noticed that T-DARPin-expressing U-CH2 cells had markedly reduced TBXT levels, which was also observed in U-CH1, JHC7, and UM-Chor1 cells (**Fig. 2a, Supplementary Fig. 2a and b**). Together with the finding of reduced *TBXT* mRNA levels (**Supplementary Fig. 2c**), this suggested that the T-DARPins altered the binding of TBXT to its super-enhancer, leading to suppression of TBXT in chordoma cells by disrupting a previously reported autoregulatory loop [5] [6]. In addition, the six T-DARPins downregulated the *YAP1* oncogene, a known transcriptional target of TBXT in chordoma (**Supplementary Fig. 2b**) [33].

**Figure 2.**
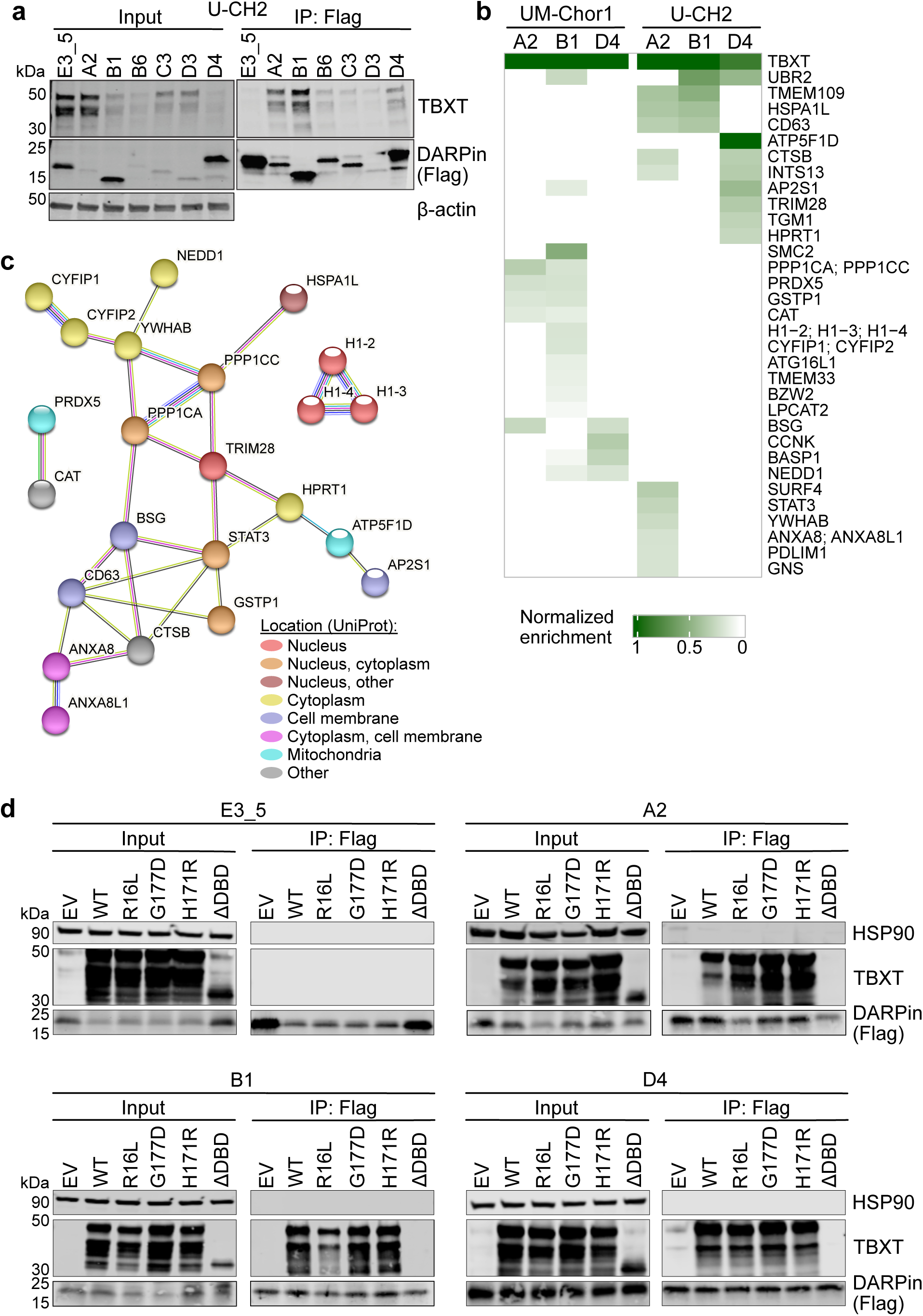
T-DARPins are specific for TBXT and bind TBXT variants. **(a)** IP of Flag-tagged DARPins from U-CH2 cells with an anti-Flag antibody five days after transduction, followed by western blotting (right). The whole-cell lysate input is shown on the left. One representative image of three independent experiments. **(b)** Enrichment heatmaps for proteins identified by AP-MS of anti-Flag IPs for DARPins A2, B1, D4, and E3_5 four days after transduction of U-CH2 and UM-Chor1 cells. **(c)** STRING network analysis with proteins detected by AP-MS. The nodes are colored according to their cellular location. The proteins detected as groups in (b) due to their peptide similarity were entered separately and are connected in the network (CYFIP1/2, ANXA8/ANXA8L1, H1-1/2/3). **(d)** Anti-Flag IPs of DARPins A2, B1, D4, and E3_5 co-transfected into HEK293T cells with EV or a vector expressing HA-tagged WT TBXT, TBXT-R16L, TBXT-G177D, TBXT-H171R, or TBXT lacking the DBD (ΔDBD).

We then focused on characterizing T-DARPins A2, B1, and D4, as they showed the strongest binding to TBXT (**Fig. 2a**) and may act via different modes as seen from the EMSA (D4 versus A2 and B1; **Fig. 1d**). To determine their selectivity, we immunoprecipitated A2, B1, D4, and E3_5 from U-CH2 and UM-Chor1 cell lysates with an anti-Flag antibody and analyzed the respective T-DARPin interactome by affinity purification mass spectrometry (AP-MS). TBXT was the only protein markedly enriched in both cell lines and with all three T-DARPins compared to E3_5 (**Fig. 2b**). Other proteins were also selectively pulled down by T-DARPins, albeit at lower levels and only in a subset of samples. Of note, we did not detect other members of the T-box transcription factor family, which comprises 17 proteins, including the founding member TBXT [34], demonstrating that T-DARPins A2, B1, and D4 are highly selective (**Fig. 2b**). For functional categorization of proteins identified by AP-MS, we used STRING (https://string-db.org) to detect protein networks and Uniprot (https://www.uniprot.org) to assign protein localization and function (**Fig. 2b, Supplementary Data 1**). Among the 33 identified proteins, 13 (including TBXT) were nuclear or nuclear membrane-associated, of which the transcription factor STAT3 and the transcriptional repressor TRIM28 were nodes in the detected STRING protein network and seemed the most promising candidate TBXT interactors due to their role in transcriptional (co-)regulation (**Fig. 2c, Supplementary Data 1**).

Single-nucleotide variants in *TBXT* have been identified in patients with sporadic chordoma and spinal developmental diseases. These include G177D in the TBXT DBD, which is a risk factor for chordoma and present in up to 94% of chordoma patients [35], and R16L and H171R, which are associated with congenital scoliosis and sacral agenesis [9, 35–37]. To investigate whether the T-DARPins bind these clinically relevant TBXT variants, we expressed HA-tagged wildtype (WT) TBXT and TBXT containing the single-nucleotide variants or lacking the DBD together with T-DARPins A2, B1, D4, or E3_5 in HEK293T cells, followed by DARPin IPs and western blotting. The three T-DARPins bound to all TBXT variants with similar efficiency than WT TBXT, whereas removing the DBD abolished DARPin binding, as expected (**Fig. 2d**). Taken together, we demonstrated the selectivity of T-DARPins for TBXT, including disease-associated TBXT variants, and identified putative TBXT-interacting proteins.

### T-DARPins impair chordoma cell proliferation, spheroid formation, and tumor growth

Next, we investigated the cellular effects of T-DARPins using chordoma cell lines. T-DARPin expression for 24 days reduced viable cells to varying degrees compared to E3_5 under regular culture conditions, with JHC7 and UM-Chor1 being the most affected and U-CH2 and U-CH12 responding the least (**Fig. 3a**). However, when we grew U-CH2 and U-CH12 cells in three-dimensional (3D) Matrigel cultures, spheroid formation was significantly reduced upon T-DARPin expression (**Fig. 3b; Supplementary Fig. 3a**) to a similar extent as when TBXT was suppressed by RNA interference (**Supplementary Fig. 3b**). The T-DARPins induced apoptosis in JHC7 but not U-CH2, UM-Chor1, and U-CH12 cells (**Fig. 3c; Supplementary Fig. 3c**). In contrast to the other cell lines, JHC7 has a genomic *TBXT* amplification [3], which may explain its higher TBXT dependence. In line with the observation that the main cellular consequence of TBXT inhibition is impaired cell proliferation, the T-DARPins increased the proportion of UM-Chor1 cells in the G0/G1 phase of the cell cycle (**Fig. 3d**). In addition, we observed morphologic changes, including loss of an organized cytoskeleton, enlarged cells with a pancake-like appearance, and narrow spindle-like cells, indicating senescence or differentiation (**Supplementary Fig. 3d**). Of note, the T-DARPins localized to the nucleus, ensuring effective TBXT inhibition in cells (**Supplementary Fig. 3d**). To verify that T-DARPins do not exert general toxic effects, we expressed them in three TBXT-negative non-chordoma cancer cell lines and did not observe reduced cell viability (**Supplementary Fig. 3e**).

**Figure 3.**
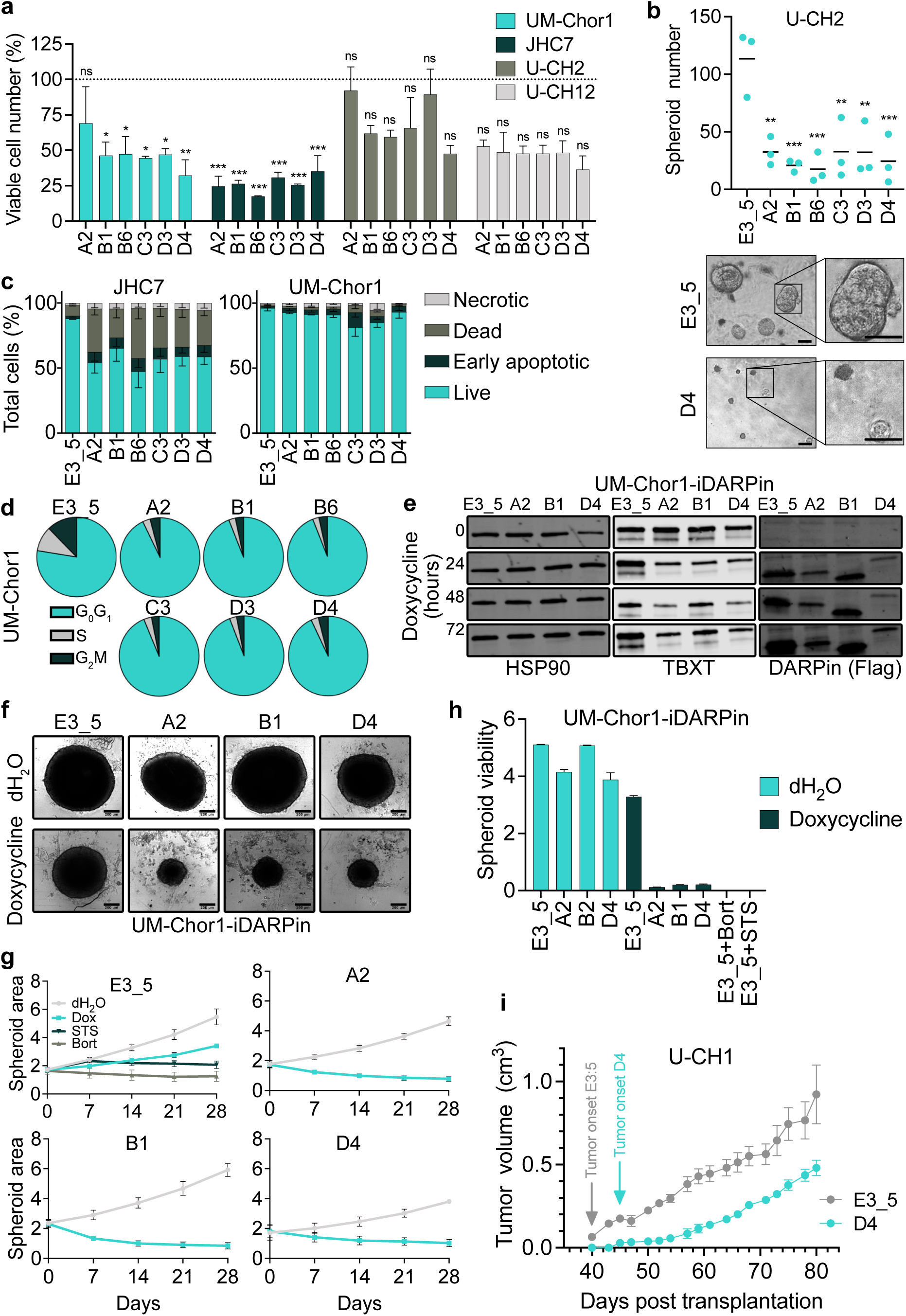
T-DARPins impair the proliferation, spheroid formation, and tumor growth of chordoma cells. **(a)** Viable cell numbers on day 24 after lentiviral transduction of chordoma cell lines with six T-DARPins relative to E3_5. One-way ANOVA with Dunnett’s test for multiple comparisons; mean + SEM of three biological replicates. *p ≤ 0.05, **p ≤ 0.01, ***p ≤ 0.001. **(b)** Spheroids of U-CH2 cells transduced with the indicated DARPins were counted after 34 days. Representative images of U-CH2 spheroids are shown below; scale bar, 50 µm. One-way ANOVA with Dunnett’s test for multiple comparisons; mean of three biological replicates. *p ≤ 0.05, **p ≤ 0.01, ***p ≤ 0.001. **(c)** Apoptosis measured by flow cytometry after staining with annexin V and propidium iodide (PI) 10 days after lentiviral transduction of JHC7 and UM-Chor1 cells with the indicated DARPins. Mean ± SEM of two biological replicates. Necrotic, PI^+^/annexinV^−^; early apoptotic, PI^−^/annexin V^+^; dead, PI^+^/annexin V^+^; live, PI^−^/annexin V^−^. **(d)** Cell cycle analysis 12 days after lentiviral transduction of UM-Chor1 cells with the indicated DARPins. One representative experiment of two biological replicates. **(e)** Western blot of UM-Chor1-iDARPin cells expressing DARPins E3_5, A2, B1, or D4 without (0 hours) or with 0.5 µg/mL doxycycline treatment for 24, 48, and 72 hours. **(f)** UM-Chor1-iDARPin spheroids imaged after 28 days without or with the addition of 0.5 µg/mL doxycycline. One representative experiment of three biological replicates. Scale bar, 200 µm. **(g)** Area (µm^2^ x 10^5^) over time of UM-Chor1-iDARPin spheroids treated with H_2_O, 0.5 µg/mL doxycycline, 10 µM staurosporine (STS), or 1 µM bortezomib (Bort). Mean ± SEM of three biological replicates with four to six spheroids averaged per treatment condition and experiment. **(h)** Viability (luminescence units x 10^7^) of UM-Chor1-iDARPin spheroids after 28 days of treatment as in (g), measured with CellTiter-Glo 3D. Mean ± SEM of two biological replicates with four to six spheroids averaged per treatment condition and experiment. **(i)** Tumor growth of U-CH1 cells transduced with T-DARPin D4 or E3_5 and injected subcutaneously into NSG mice. Shown is the mean volume ± SEM of six tumors per condition until the first mouse had to be sacrificed because the maximum allowable tumor size was reached.

The greater TBXT dependence of chordoma cells in 3D cultures is consistent with TBXT’s function in tissue organization during development [38]. Since faithful chordoma models for biology research and drug discovery are needed [11], we established UM-Chor1 spheroids using ultra-low attachment plates and confirmed that they are suited for long-term culture by detecting stable TBXT expression and proliferation, as determined by Ki-67 positivity, after 45 days (**Supplementary Fig. 3f**). To enable fast induction of T-DARPins and subsequent TBXT inhibition in spheroids, we engineered UM-Chor1 cells, named UM-Chor1-iDARPin cells hereafter, to express the tetracycline-responsive transactivator rtTA3 and contain a CMV promoter with multiple doxycycline-responsive elements that drive DARPin expression. We observed tight control of DARPin induction with no expression in the absence of doxycycline and rapid expression of E3_5 and T-DARPins A2, B1, and D4 in the presence of doxycycline (**Fig. 3e**). The downregulation of TBXT protein observed after 12 days with lentiviral DARPin delivery (**Supplementary Fig. 2a and b**) was seen as early as 24 hours in UM-Chor1-iDARPin cells, suggesting rapid interruption of the TBXT autoregulatory loop (**Fig. 3e**). Induction of T-DARPins A2, B1, and D4 in seven-day-old UM-Chor1-iDARPin spheroids resulted in complete growth arrest over 28 days, similar to treatment with the broad-spectrum kinase inhibitor staurosporine or the proteasome inhibitor bortezomib, as evidenced by smaller spheroid size and reduced cell viability (**Fig. 3f–h**).

To determine the efficacy of T-DARPins *in vivo*, we injected U-CH1 cells stably expressing E3_5 or T-DARPin D4 subcutaneously into NOD.Cg-*Prkdc^scid^ Il2rg^tm1Wjl^*/SzJ (NSG) mice (**Supplementary Fig. 3g**). Cells had robust DARPin expression and reduced TBXT levels with T-DARPin D4 before injection (**Supplementary Fig. 3h**). U-CH1 E3_5 yielded tumors at all six injection sites after 40 days, while the appearance of the first U-CH1 D4 tumor was delayed by one week, and tumor growth was slowed in the first three weeks compared to E3_5 tumors (**Fig. 3i**), demonstrating the capacity of T-DARPins to inhibit chordoma growth *in vivo*. However, the tumors then grew at similar rates as the control tumors, suggesting the outgrowth of cells that had developed resistance to TBXT inhibition (**Fig. 3i**). Consistent with this, D4 tumors had lost DARPin expression and restored TBXT expression at the end of the experiment (**Supplementary Fig. 3i**). PCR amplification of DARPin transcripts from the tumors revealed that D4 contained a deletion of approximately 100 base pairs, whereas the length of E3_5 transcripts remained unaffected (**Supplementary Fig. 3j**). Together, these results demonstrated that T-DARPins effectively inhibit endogenous TBXT, resulting in reduced proliferation and spheroid growth of chordoma cells and impaired tumor formation in xenografts.

### T-DARPins cause widespread transcriptomic and proteomic changes

Despite the knowledge that TBXT is the essential chordoma driver, the TBXT-regulated transcriptome is not fully understood. We performed RNA sequencing (RNA-seq) of UM-Chor1 cells transduced with E3_5 or T-DARPin A2, B1, or D4. The changes induced by the three T-DARPins relative to E3_5 were highly concordant (**Supplementary Fig. 4a; Supplementary Data 2**). We found 1,803 significantly downregulated and 1,468 significantly upregulated genes (log_2_(fold change) > 0.585 or < −0.585 [corresponding to a 1.5-fold change], false discovery rate [FDR] < 1%) that were common to the three T-DARPins (**Fig. 4a; Supplementary Data 2**), demonstrating that TBXT has widespread transcriptional properties in chordoma cells.

**Figure 4.**
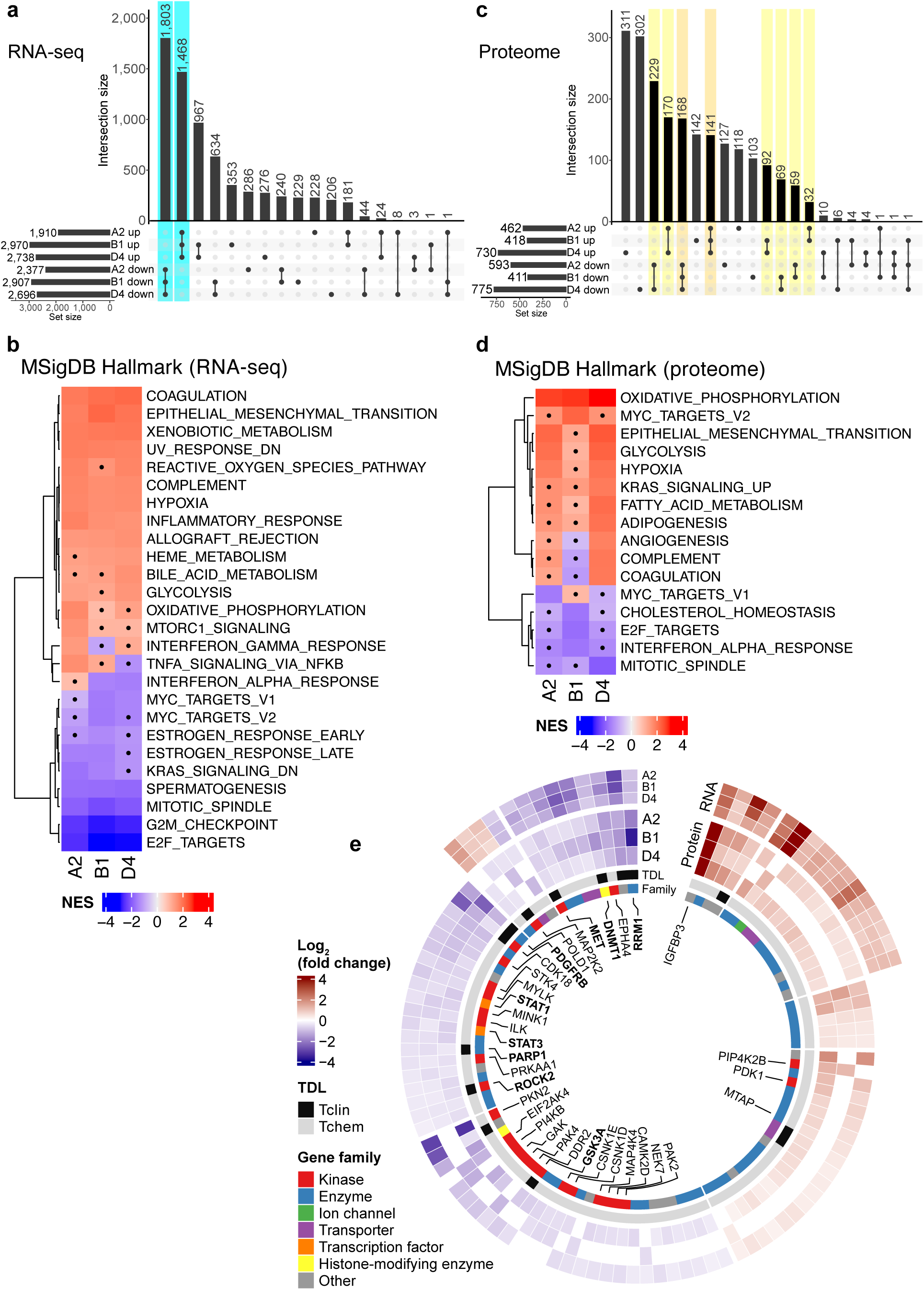
Assessment of the TBXT regulome by T-DARPin expression in UM-Chor1 cells. **(a)** UpSet plot of RNA-seq data showing the number and intersections of genes significantly deregulated by T-DARPin expression as a matrix of six sets (T-DARPins A2, B1, and D4; up- or downregulated). Rows show the number of genes per set. Columns show the number of intersecting genes at the top and the intersecting sets as connected dots at the bottom. DEGs, as defined in the text, are indicated in teal. **(b)** GSEA of RNA-seq data with the MSigDB Hallmark gene collection. NES, normalized enrichment score. **(c)** UpSet plot of DDA-MS data showing the number and intersections of proteins significantly deregulated by T-DARPin expression as a matrix of six sets (T-DARPins A2, B1, and D4; up- or downregulated). Rows show the number of proteins per set. Columns show the number of intersecting proteins at the top and the intersecting sets as connected dots at the bottom. DEPs, as defined in the text, are indicated in orange and yellow. **(d)** GSEA of DDA-MS data with the MSigDB Hallmark gene collection. NES, normalized enrichment score. **(e)** Circos plot of DEPs categorized according to their target development levels (TDL). The inner heatmap shows the log_2_(fold change) of proteins deregulated by T-DARPins A2, B1, and D4. The outer heatmap shows the log_2_(fold change) of the corresponding RNA-seq data if the respective DEP-encoding gene was also among the DEGs.

Among the 3,271 differentially expressed genes (DEGs) were TBXT targets previously identified by non-protein-based methods such as CRISPR knockout or degron tagging [5, 6], further validating the on-target activity of T-DARPins. These included the embryonic notochord markers *ACAN*, *COL2A1*, *SOX9*, and *TBXT* itself, which are abnormally active in chordoma cells [39]; *TPX2*, *C7orf69*, *WEE1*, *ATP6V1B2*, and *SAE1*, which are controlled by binding of TBXT to their super-enhancers [5, 6]; collagen genes associated with the embryonic notochord (*COL17A1*, *COL26A1*) [40] or chordoma (*COL11A2*) [41]; and the chordoma-associated cytokeratin genes *KRT18* and *KRT19* [42, 43] (**Supplementary Fig. 4b and c**). Gene set enrichment analysis (GSEA) using the Hallmark, ARCHS^4^ Tissue, and Gene Ontology Biological Process (GO_BP) gene signature collections (**Fig. 4b, Supplementary Fig. 4d-f, Supplementary Data 3**) revealed significant suppression of networks related to cell cycle checkpoint regulation, cell division, and DNA replication and repair (**Fig. 4b, Supplementary Fig. 4e**) and of gene signatures associated with embryonic stem and progenitor cells, most likely reflecting TBXT-regulated gene programs of remnant notochordal cells (**Supplementary Fig. 4d**). Gene signatures upregulated upon TBXT inhibition were mainly associated with differentiated cell types, e.g., of muscle or neuronal origin, and epithelial-mesenchymal transition (**Fig. 4b, Supplementary Fig. 4d**), hinting towards differentiation of chordoma cells after TBXT inhibition, and with various metabolic processes and inflammatory response (**Fig. 4b, Supplementary Fig. 4f**).

We next used data-dependent acquisition mass spectrometry (DDA-MS) to determine the TBXT-dependent proteome in UM-Chor1 cells following expression of T-DARPins A2, B1, or D4 for 14 days or CRISPR/Cas9-mediated TBXT knockout (**Supplementary Figure 4g and h**). We identified 960 significantly differentially expressed proteins (DEPs; log_2_(fold change) > 0.2 or < −0.2 [corresponding to a 1.15-fold change], FDR < 5%) associated with at least two T-DARPins and regulated in the same direction, of which 525 were downregulated and 435 were upregulated (**Fig. 4c; Supplementary Figure 4i; Supplementary Data 4**). Of these, 523 were also significantly deregulated in the same direction by TBXT knockout (256 downregulated and 267 upregulated; **Supplementary Data 4**). Most prominent was the extreme upregulation of insulin-like growth factor binding protein 3 (IGFBP3), which was present in all four conditions (**Supplementary Fig. 4i**). GSEA of T-DARPin proteomes using the same gene signature collections as for the RNA-seq analysis (**Fig. 4d, Supplementary Fig. 4j–l, Supplementary Data 5**) revealed suppression of networks involved in cell cycle regulation, DNA replication and repair, and interferon alpha response after TBXT inhibition (**Fig. 4d, Supplementary Fig. 4k**) and upregulation of signatures associated with differentiated cell types, epithelial-mesenchymal transition, and metabolic processes, particularly oxidative phosphorylation and glycolysis (**Fig. 4d, Supplementary Fig. 4j and l**), which was largely consistent with the result of GSEA of T-DARPin transcriptomes. Taken together, TBXT inhibition caused significant changes in the transcriptome and proteome of chordoma cells indicative of impaired cell division, a switch from an embryonic to a more differentiated cell state, an interferon/inflammatory response, the induction of various metabolic processes, and perturbed DNA replication and repair.

### The TBXT regulome includes potential pharmacologic targets

In addition to TBXT itself, essential downstream effectors may also represent therapeutic targets that could be addressed alone or in combination with future TBXT-directed drugs. To identify such candidate drug targets, we used The Cancer Druggable Gene Atlas [44] and categorized the TBXT-regulated DEGs and DEPs according to the target development levels Tclin (clinically approved drugs available), Tchem (experimental drugs available), Tbio (potentially druggable), and Tdark (currently not druggable). Overall, 1,065 DEGs and/or DEPs could be assigned to any of the four categories, of which 102 met the Tclin level and 291 the Tchem level (**Supplementary Data 6**). Tclin and Tchem targets were identified in both the transcriptome and proteome (n = 6 and n = 23, respectively), the proteome only (n = 8 and n = 63, respectively), and the transcriptome only (n = 88 and n = 205, respectively; **Supplementary Fig. 4m, Supplementary Data 6**). As proteins are the direct targets of drugs, we subsequently focused on DEPs with Tclin (n = 14) and Tchem (n = 86) levels and not on DEGs (**Fig. 4e**). However, the large group of candidates identified only by RNA-seq may include relevant targets, such as the clinically actionable kinases RET, NTRK1, and CDK6 (**Supplementary Fig. 4m**).

Sixty of the 100 Tclin/Tchem DEPs were suppressed by TBXT inhibition and, therefore, represent effectors directly or indirectly elevated by TBXT. Among these were 24 kinases, e.g., MET, GSK3A, ROCK2, and PDGFRB; the enzymes RRM1 and PARP1; and the DNA methyltransferase DNMT1 (**Fig. 4e, Supplementary Data 6**). While PDGFRB and PARP1 inhibitors have been applied as chordoma therapies [14, 45], the biological and clinical role of the other candidates is unclear. The receptor tyrosine kinase MET and its ligand hepatocyte growth factor are strongly expressed in primary chordoma samples, and chordoma cell lines, particularly of sacral origin, are sensitive to MET inhibition, most efficiently together with EGFR inhibition [13, 46]. RRM1, the large catalytic subunit of ribonucleotide reductase responsible for generating deoxynucleoside triphosphates for DNA synthesis, is inhibited by gemcitabine [47], which is used to treat various cancers but whose efficacy in chordoma has not yet been tested. Similarly, the catalytic activity of DNMT1 can be blocked by azacitidine or decitabine, both used to treat acute myeloid leukemia. Finally, DEPs positively regulated by TBXT also included the transcription factors STAT1 and STAT3, which can be modulated by inhibiting upstream regulatory JAK kinases, for which several approved drugs are available.

Forty Tclin/Tchem DEPs were upregulated by TBXT inhibition and are thus directly or indirectly suppressed by TBXT in chordoma cells. Among these was the 5-methylthioadenosine phosphorylase MTAP (**Fig. 4e**). The *MTAP* gene is co-deleted with the tumor suppressor gene *CDKN2A* in multiple cancer types, causing sensitivity to inhibition of the protein methyltransferase PRMT5 [48, 49], and several PRMT5 inhibitors are being tested in clinical trials. It is, therefore, tempting to speculate that TBXT-mediated MTAP suppression may phenocopy this effect in chordoma. In addition, several cathepsins (CTSA, CTSB, and CTSL) were among the Tchem proteins induced by TBXT inhibition. Cathepsins are frequently implicated in cancer development and represent emerging drug targets [50, 51]. Finally, the most upregulated candidate in both the proteome and the transcriptome was IGFBP3, which belongs to the Tchem category. In summary, the TBXT regulome contains multiple pharmacologically tractable proteins, including some targets of clinically approved drugs, and therefore represents a valuable resource for future chordoma drug discovery campaigns.

### The TBXT regulome is partially modulated by IGFBP3

IGFBP3 is a signaling molecule implicated in cell survival and apoptosis [52], depending on cellular context [53, 54]. It is secreted and glycosylated [55–57] and exerts para- and autocrine effects [58]. To study the role of IGFBP3 in chordoma, we first confirmed its upregulation using ELISA in UM-Chor1 cells after TBXT knockout (**Fig. 5a**) and in JHC7 and U-CH2 cells after TBXT inhibition with T-DARPins (**Supplementary Fig. 5a**). IGFBP3 was also detected in conditioned medium of UM-Chor1 and U-CH2 cells, in particular after TBXT loss, and treatment with the glycosidase PNGase F decreased the molecular weight of IGFBP3 (**Fig. 5b, Supplementary Fig. 5b**). This indicated that IGFBP3 is secreted and glycosylated in chordoma cells, particularly evident after TBXT loss. Since IGFBP3 is a YAP1 target [59], and YAP1 is transcriptionally regulated by TBXT [33], we examined whether IGFBP3 induction upon TBXT inhibition is mediated via YAP1. sgRNA knockout or shRNA knockdown of YAP1 in UM-Chor1 cells did not affect IGFBP3 mRNA and protein expression, whereas TBXT knockout increased *IGFBP3* mRNA as expected (**Fig. 5c, Supplementary Fig. 5c and d**), demonstrating that the reciprocal relationship between TBXT and IGFBP3 expression is independent of YAP1.

**Figure 5.**
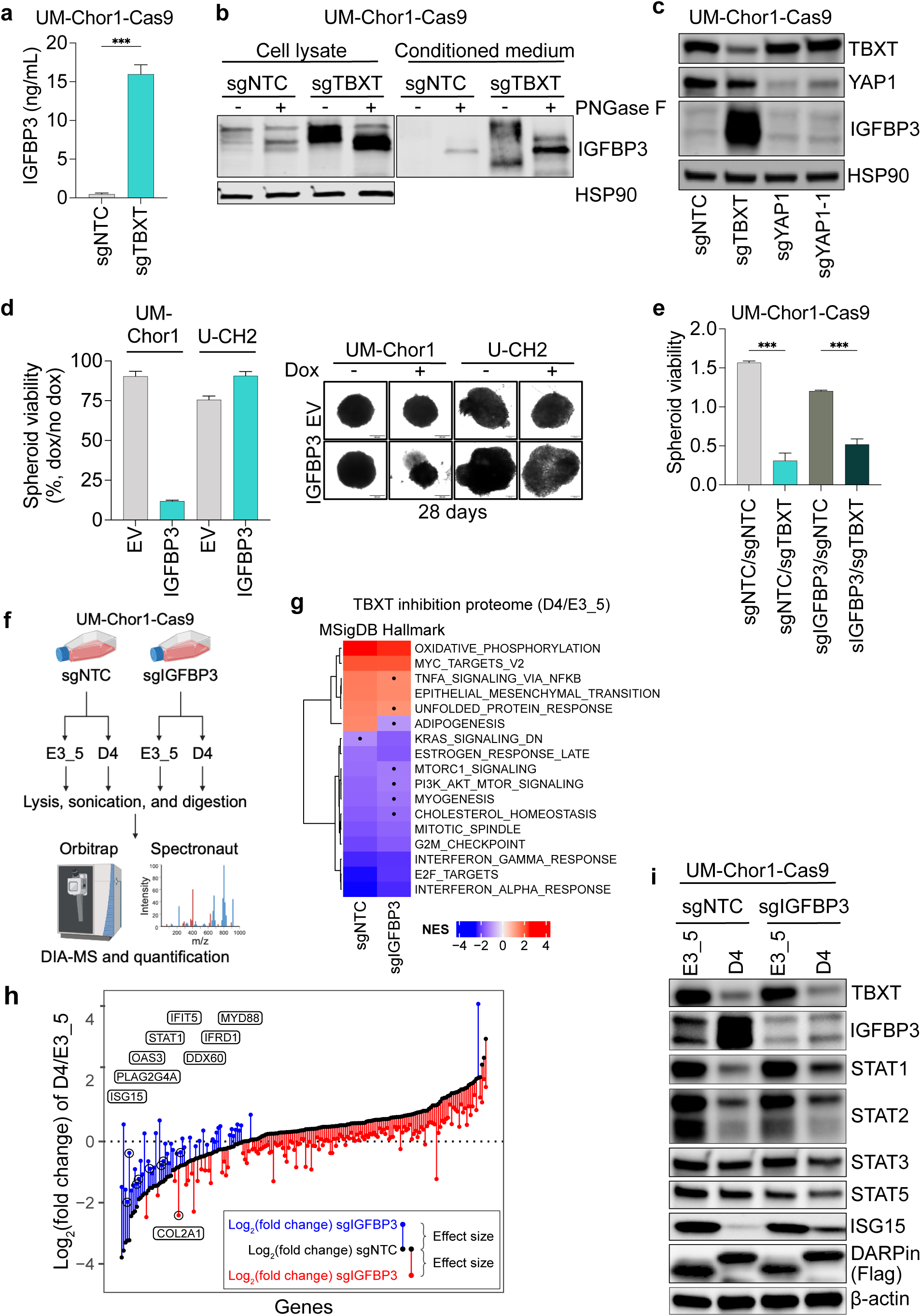
IGFBP3 modifies part of the TBXT regulome. **(a)** IGFBP3 protein levels quantified by ELISA of UM-Chor1-Cas9 cells transduced with sgNTC or sgTBXT. Mean + SEM of three biological replicates. Unpaired t-test. ***p ≤ 0.001. **(b)** IGFBP3 protein expression in lysates and culture medium of UM-Chor1-Cas9 cells transduced with sgNTC or sgTBXT. Samples were treated with or without the glycosidase PNGase F. **(c)** Western blot of UM-Chor1-Cas9 cells transduced with sgNTC, sgTBXT, or two sgRNAs targeting YAP1. **(d)** Viability of UM-Chor1 and U-CH2 spheroids stably expressing doxycycline-inducible IGFBP3 or EV and treated with 0.5 µg/mL doxycycline compared to untreated spheroids after 28 days. Mean + SEM of two biological replicates with five to six spheroids averaged per treatment condition and experiment. Representative images of single spheroids are shown on the right. **(e)** Viability (luminescence units x 10^7^) after 14 days of UM-Chor1-Cas9 spheroids double-transduced with sgNTC/sgNTC, sgTBXT/sgNTC, sgIGFBP3/sgNTC, or sgIGFBP3/sgTBXT. Mean + SEM of three biological replicates with four to six spheroids averaged per treatment condition and experiment. One-way ANOVA with Dunnett’s test for multiple comparisons; ***p ≤ 0.001. **(f)** Schematic of the DIA-MS analysis of UM-Chor1-Cas9 cells 14 days after transduction with sgNTC or sgIGFBP3 and subsequently with T-DARPin D4 or E3_5. Created in BioRender. Fröhling, S. (2024) BioRender.com/r29p687. **(g)** MSigDB Hallmark GSEA comparing the proteomes of sgNTC and sgIGFBP3 UM-Chor1-Cas9 cells in the context of TBXT inhibition. **(h)** Expression changes of IGFBP3-dependent genes induced by T-DARPin D4 in the context of transduction with sgNTC (black dots) or sgIGFBP3 (red and blue dots). Differences between log_2_(fold changes) are depicted by vertical lines between dots. Red dots and lines show genes whose log_2_(fold change) decreased with IGFBP3 knockout. Blue dots and lines show genes whose log_2_(fold change) increased with IGFBP3 knockout. Genes involved in the interferon response pathway are indicated above the curve, and the corresponding data points are circled. **(i)** Western blot of UM-Chor-Cas9 cells with (sgIGFBP3) or without (sgNTC) IGFBP3 knockout and expressing DARPins E3_5 or D4.

We next hypothesized that high IGFBP3 levels might be toxic to chordoma cells and, therefore, suppressed by TBXT and that the proliferation arrest caused by TBXT inhibition might be mediated by IGFBP3 re-expression. Consistent with this, doxycycline-induced IGFBP3 overexpression decreased the viability and size of UM-Chor1 spheroids after 28 days. In contrast, U-CH2 spheroids were unaffected (**Fig. 5d**), suggesting a cell line- and chordoma type-specific effect of IGFBP3. Additional depletion of IGFBP3 partially restored TBXT knockout-induced UM-Chor1 spheroid viability (**Fig. 5e**), indicating that IGFBP3 upregulation contributes to the growth-inhibitory effect of TBXT inhibition.

To investigate the influence of IGFBP3 on the TBXT regulome at high resolution, we performed data-independent acquisition mass spectrometry (DIA-MS) in UM-Chor1 cells stably transduced with an sgRNA targeting IGFBP3 (sgIGFBP3) or a non-targeting control sgRNA (sgNTC) in combination with T-DARPin D4 or the control DARPin E3_5 (**Fig. 5f, Supplementary Data 7**). In sgNTC cells (no IGFBP3 knockout), 1,752 proteins were upregulated, and 1,749 proteins were downregulated by TBXT inhibition via D4 relative to E3_5 (log_2_(fold change) > 0.2 or < −0.2 [corresponding to a 1.15-fold change], FDR < 5%). With IGFBP3 knockout, fewer proteins were deregulated by TBXT inhibition (1,373 up and 1,586 down), of which 81% overlapped with the proteins deregulated in sgNTC cells (**Supplementary Fig. 5e**). GSEA with proteins regulated by TBXT in unperturbed compared to IGFBP3 knockout cells using the Hallmark gene signature collection revealed enrichment of oxidative phosphorylation and epithelial-mesenchymal transition and suppression of cell cycle-related networks and interferon response (**Fig. 5g**), confirming the effects of TBXT inhibition found by DDA-MS (**Fig. 4d**).

To determine the proteins whose levels are influenced by TBXT activity and IGFBP3 expression, we analyzed the DIA-MS data by multifactorial linear regression with interactions. This identified 193 proteins, of which 144 (75%) were suppressed and 49 (25%) were upregulated after IGFBP3 loss (**Fig. 5h**). Notably, most of the proteins downregulated by IGFBP3 loss were upregulated by TBXT inhibition and vice versa, suggesting that IGFBP3 mainly counteracts the regulatory effects of TBXT (**Fig. 5h**). Among the modulated proteins was the embryonic notochord marker COL2A1, whose TBXT inhibition-induced suppression was further enhanced by IGFBP3 loss, whereas other notochord markers, including TBXT itself, were unaffected by IGFBP3 (**Fig. 5h, Supplementary Fig. 5f**). Eight modulated proteins are involved in the interferon response pathway, including STAT1 and the interferon-stimulated genes (ISGs) ISG15, OAS3, IFRD1, and IFIT5, whose expression was downregulated by TBXT inhibition and partially restored by IGFBP3 loss (**Fig. 5h**). Western blot analysis showed that TBXT inhibition strongly downregulated STAT1, STAT2, and ISG15 in UM-Chor1 cells, which was partially reversed in the absence of IGFBP3 in the case of STAT1 and ISG15 but not STAT2, confirming the proteomics results (**Fig. 5i**). For completeness, we also blotted for STAT3 and STAT5 and found a slight decrease in both proteins upon TBXT inhibition which was enhanced in the absence of IGFBP3.

Finally, since pathway enrichment analysis showed little difference between TBXT-inhibited proteomes with (sgIGFBP3) or without (sgNTC) IGFBP3 depletion, we explored differentially enriched pathways by ranking proteins with the moderated t-statistic upon multifactorial analysis and applying GSEA with the MSigDB Hallmark gene set. Oxidative phosphorylation, epithelial-mesenchymal transition, and the interferon alpha and gamma response pathways were the only significantly enriched gene sets (**Supplementary Fig. 5g**). This suggested that although multiple pathways respond to TBXT inhibition, these four may be particularly influenced by both TBXT and IGFBP3. Together, we showed that part of the TBXT regulome is modulated by IGFBP3, particularly factors involved in the interferon response, suggesting that IGFBP3 can fine-tune the action of TBXT to precisely fulfill the requirements of chordoma cells for survival and proliferation.

### TBXT promotes JAK-STAT signaling and renders chordoma cells sensitive to JAK2 inhibition

The multi-omics experiments described above pointed to a functional link between TBXT and interferon signaling, particularly the JAK-STAT axis. Specifically, we detected STAT3 as a potential TBXT binding partner (**Fig. 2b**), found interferon response signaling to be significantly affected by TBXT inhibition on the transcriptomic (**Fig. 4b**) and proteomic (**Fig. 4d and 5g, Supplementary Fig. 5g**) level, and observed downregulation of STATs and interferon-stimulated genes after TBXT inhibition (**Fig. 5h and i**). To substantiate this association, we mapped the levels of ISGs across the RNA-seq and proteomics experiments and observed downregulation of ISGs upon TBXT inhibition in all data sets (**Fig. 6a**), suggesting that chordoma cells might have high baseline interferon signaling driven by TBXT. To ensure that this was not only a feature of chordoma cell lines, we analyzed RNA-seq data of 1,077 tumor samples from 1,008 cancer patients, including 12 samples from nine chordoma patients, enrolled in DKFZ/NCT/DKTK MASTER, a cross-entity precision oncology program focusing on rare cancers [60]. In line with the observations in cell lines, MSigDB enrichment scores revealed interferon alpha and gamma response as the most enriched signaling axes in chordoma but not in the pan-cancer cohort (**Fig. 6b**).

**Figure 6.**
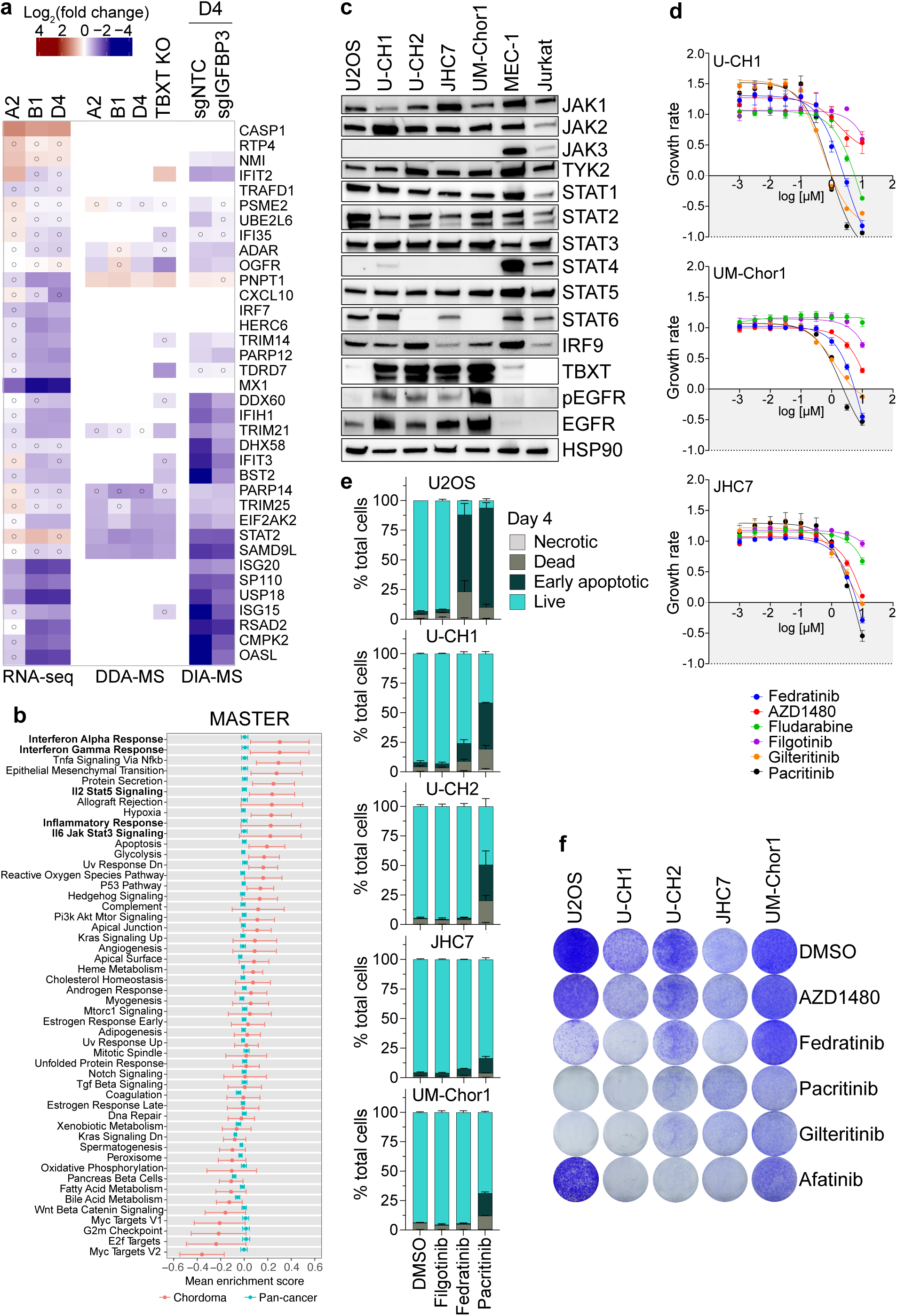
Chordoma cells depend on JAK-STAT signaling. **(a)** ISG expression in UM-Chor1 cells as determined by RNA-seq, DDA-MS, and DIA-MS. The different treatment conditions are indicated at the top. Open circles indicate that expression was detected but was not significant. **(b)** GSEA of 1,077 tumor transcriptomes from 1,008 patients using 50 Hallmark gene sets. The mean gene set enrichment scores of 12 chordoma samples (red dots) were plotted along with those of 1,065 non-chordoma samples (blue dots), which represent the cohort average for each gene set. Gene sets were ordered from the highest to lowest mean enrichment score in the chordoma group. Error bars represent 95% confidence intervals. **(c)** Western blots of chordoma cell lines U-CH1, U-CH2, JHC7, and UM-Chor1, the osteosarcoma cell line U2OS, the B-cell leukemia cell line MEC-1, and the T-cell leukemia cell line Jurkat to detect expression of JAK/STAT pathway members, TBXT, EGFR, and phospho-EGFR (Y1068). **(d)** GR analysis of the indicated cell lines after seven days of treatment with the pan-JAK inhibitor filgotinib, the JAK2 inhibitors AZD1480, fedratinib, and pacritinib, the STAT1 inhibitor fludarabine, and the FLT3 inhibitor gilteritinib at concentrations of 10 µM to 1 nM. GR values between 1 and 0 indicate proliferation inhibition, 0 indicates a complete cytostatic effect, and values between 0 and −1 indicate additional cytotoxicity. One of three representative experiments is shown, with each data point representing a technical triplicate. **(e)** Apoptosis measured by flow cytometry after staining with annexin V and 7-AAD of cell lines treated with 3 µM of the indicated drugs for four days. Mean + SEM of two biological replicates. **(f)** Colony formation of U2OS and chordoma cell lines treated with 1 µM of the indicated drugs for 14 days. Representative images of three independent experiments (**Supplementary Fig. 6d**).

To better understand JAK-STAT activity in chordoma, we examined the levels of JAK and STAT family proteins in multiple chordoma cell lines, the osteosarcoma cell line U2OS, the chronic B-cell leukemia cell line MEC-1, and the T-cell leukemia cell line Jurkat. U2OS was selected as a non-chordoma sarcoma cell line sensitive to JAK inhibition [61], and MEC-1 and Jurkat as lymphoid cells with intrinsically high JAK/STAT signaling. As expected, all chordoma cell lines lacked JAK3, which is only expressed in hematopoietic lineages [62]. They also lacked STAT4 but expressed JAK1, JAK2, and TYK2 to varying degrees, as well as STAT1, STAT2, STAT3, and STAT5 to similar levels as MEC-1 and Jurkat (**Fig. 6c**). STAT6 was expressed in U-CH1 and JHC7 cells but not UM-Chor1 or U-CH2 cells (**Fig. 6c**). In addition, all chordoma cell lines expressed IRF9, which forms a complex with activated STAT1 and STAT2 that stimulates the transcription of ISGs [62].

Based on these results, we reasoned that chordoma cells might be dependent on members of the JAK-STAT pathway, representing a novel vulnerability that can be addressed by clinically approved drugs. To test this, we measured the sensitivity of chordoma cells to the pan-JAK inhibitor filgotinib [63]; the JAK2 inhibitors AZD1480 [64], fedratinib [65], and pacritinib [66]; the chemotherapeutic and STAT1 inhibitor fludarabine [67]; and the FLT3 inhibitor gilteritinib as a control [68], since fedratinib and pacritinib also inhibit FLT3. JAK-dependent U2OS cells were used as a positive control. Drug responses were calculated using growth rate (GR) analysis [69] to account for the highly variable cell division rates of chordoma cell lines. While fludarabine, AZD1480, and filgotinib were mainly cytostatic, the JAK2 inhibitors fedratinib and pacritinib had the strongest effect on all cell lines and pushed the GR below 0, indicating a cytotoxic effect (**Fig. 6d, Supplementary Fig. 6a**). Unexpectedly, gilteritinib was equally toxic; however, since chordoma cells do not express FLT3 (**Supplementary Fig. 6b**), this was probably a non-specific effect.

Consistent with their predicted cytotoxicity, fedratinib and pacritinib, but not the pan-JAK inhibitor filgotinib, induced apoptosis in U-CH1 cells after four days, which increased after seven days (**Fig. 6e, Supplementary Fig. 6c**). Additionally, pacritinib induced apoptosis in U-CH2, UM-Chor1, and JHC7 cells albeit to a lesser degree. Furthermore, we observed markedly reduced colony formation over 14 days after JAK2 inhibition in U2OS and chordoma cells, particularly U-CH1, with pacritinib being more effective than fedratinib (**Fig. 6f, Supplementary Fig. 6d**). The EGFR inhibitor afatinib, which we included as a positive control since it downregulates TBXT [16, 22], was equally harmful to chordoma cells but, as expected, not to the TBXT-negative and EGFR-inactive U2OS cells (**Fig. 6c**). The FLT3 inhibitor gilteritinib also strongly inhibited colony formation. In summary, our data indicate that chordoma cells require JAK-STAT signaling driven by TBXT, in particular JAK2, suggesting an immediately applicable strategy for targeted treatment of chordoma patients using clinically approved JAK2 inhibitors.

## DISCUSSION

TBXT plays a crucial role in regulating chordoma cell function. Consequently, it is an attractive target for developing anti-chordoma drugs, which should provide a substantial therapeutic window due to the largely absent TBXT expression in normal tissues. However, many transcription factors, like TBXT, evade pharmacologic targeting. Thus, new approaches are needed to find targeted therapies for transcription factor-driven cancers. We developed DARPins that effectively and selectively bind TBXT and inhibit its transcriptional function in chordoma cells. Using these novel agents, we determined the TBXT regulome and uncovered components of TBXT-dependent signaling networks that may represent therapeutic vulnerabilities.

We developed T-DARPins for two reasons: to generate tools for studying the function of TBXT in chordoma or other TBXT-expressing cancers in a highly specific manner and to provide the basis for molecular mechanism-instructed therapies. T-DARPins A2, B1, and D4 were highly concordant in blocking the transcriptional activity of TBXT by impairing its binding to DNA and rapidly downregulating TBXT mRNA and protein. While binding to clinically relevant TBXT variants associated with chordoma and developmental disorders of the axial skeleton was retained, the T-DARPins were highly selective, as they did not bind T-box homologs of TBXT.

So far, DARPins have been developed against secreted, cell surface, and cytoplasmic targets [31, 70, 71], and some molecules have entered clinical trials [28, 72]. Our T-DARPins demonstrate that nuclear proteins can also be bound and blocked by DARPins without adding a nuclear localization signal. Based on these properties, T-DARPins would, in principle, be suitable as therapeutic agents. However, the delivery into tumor cells is challenging for further clinical development. Potential strategies include fusion with bacterial toxins that allow translocation from the endosome to the cytosol, as demonstrated in cell culture [70, 71, 73], fusion with cell-penetrating-peptides [74, 75], fusion with domains forming a complex with cationic and ionizable lipids via electrostatic interactions [76], mRNA-based approaches [77], and viral systems, e.g., shielded adenoviruses that minimize targeting in the liver and recognition by the immune system [30, 78]. In addition to a potential role as clinically applied biologics, T-DARPins could also serve as a starting point for developing TBXT-directed small molecules.

As expected, due to the essential role of TBXT in chordoma biology [3, 5, 6, 16, 17, 33, 35, 39] we observed impaired cell growth by T-DARPin-mediated TBXT inhibition in various experimental models. This effect was stronger in 3D models, highlighting the importance of using such formats for chordoma drug discovery, as also demonstrated in a recent proof-of-concept study testing drug sensitivity in patient-derived chordoma organoids [11]. In addition, T-DARPin D4 delayed tumor onset and initial growth in a U-CH1 xenograft. The finding that surviving U-CH1 cells lost T-DARPin expression by acquiring a deletion in the D4 transgene illustrated the selection pressure to restore TBXT.

The predominant mechanism by which T-DARPins impaired chordoma cell growth was cell cycle disruption, as evidenced by G0/G1 arrest and the most enriched signaling networks in the TBXT-regulated transcriptome and proteome. The effect on the cell cycle and the morphologic changes after TBXT inhibition suggest that chordoma cells undergo senescence – a fate that has been previously reported [3, 6]. One chordoma cell line, JHC7, underwent apoptosis upon T-DARPin expression. This exception may be related to genomic *TBXT* amplification in these cells [3], which could be associated with particularly strong TBXT dependence.

In addition, some UM-Chor1 cells assumed an elongated, neuron-like shape after T-DARPin expression, indicating differentiation [3]. This was also supported by the transition from transcriptional programs resembling those of embryonic stem cells to those of differentiated cell types. Since TBXT is an essential driver of mesoderm formation and notochord development [79], its continued expression and transcriptional activity in chordoma leads to the persistence of an immature cellular phenotype [6]. Consequently, TBXT inhibition could impose a more differentiated identity on chordoma cells. UM-Chor-1 cells seemed to be driven towards the neural lineage. However, additional studies with multiple chordoma cell lines or primary chordoma organoids are needed to investigate all possible fates of chordoma cells after TBXT inhibition and to develop biomarkers of differentiation for guiding future TBXT-targeted therapies.

Taking into account the transcripts and proteins deregulated by three independent T-DARPins allowed us to confidently map the TBXT regulome of UM-Chor-1 chordoma cells. Besides the main impact of TBXT on the cell cycle and identity of chordoma cells, analysis of the core TBXT regulome identified additional gene programs. In particular, glycolysis and oxidative phosphorylation were upregulated after TBXT blockade in both the transcriptome and proteome. Two previous studies have also suggested a role for metabolic processes in chordoma. Using CRISPR/Cas9 screening in chordoma cell lines, Sharifnia et al. [80] identified essential genes related to glucose, amino acid, and cholesterol metabolism. Proteomic analysis of biopsies from chordoma patients revealed that oxidative phosphorylation was strongly deregulated [81]. Further studies are needed to determine the exact role of these pathways in chordoma biology and their potential for clinical exploitation.

By assigning target development levels to all genes and proteins in the TBXT regulome, we identified candidates that could be exploited as therapeutic entry points, alone or in combination with future anti-TBXT treatments to increase efficacy. This strategy recovered known targets of small-molecule inhibitors, including PDGFRB, CDK6, and PARP1, that have been [14, 45] or are being (ClinicalTrials.gov: NCT03110744 and NCT03127215) tested in clinical trials enrolling chordoma patients, demonstrating the potential to identify druggable molecules among the TBXT-regulated proteins. Inhibition of MET, whose anti-chordoma activity has been shown preclinically [46], DNMT1 by repurposing drugs used in acute myeloid leukemia, and PRMT5 in the context of TBXT-induced MTAP suppression [48, 49], as well as treatment with cytotoxic drugs that inhibit RRM1 [47], such as gemcitabine, appear to be the most promising in this respect.

In addition to immediately clinically actionable targets, the TBXT regulome contained many potentially druggable candidates. IGFBP3 stood out for its extreme upregulation after TBXT loss in several chordoma cell lines. Of note, IGFBP3 is also upregulated in Ewing sarcoma cells after suppression of the EWS::FLI1 fusion [82], indicating that it may be a more common downstream effector of oncogenic transcription factors in mesenchymal malignancies. IGFBP3 is an intra- and extracellular signaling protein with pro- and antitumor activity. It displaces IGF1 from its receptor, thereby inhibiting the IGF1-IGF1R signaling cascade and reducing cell growth [83]. However, IGFBP3 also has been proposed to have IGF-independent functions, potentially explaining its pleiotropic effects in different cellular contexts. For example, it interacts with TGF-β signaling, binds to its own receptor mediating apoptosis, is suspected to be transported into the nucleus via importin-β to act as a transcription factor or transcriptional co-regulator, where it could influence many cellular processes, including DNA repair, and can be cleaved by metalloproteases, with the resulting fragments having diverse functions [83]. We found that IGFBP3 is glycosylated and secreted in chordoma cells, especially when it is highly expressed after TBXT inhibition. Our initial data on IGFBP3 function in chordoma cells showed that it impairs spheroid growth and that its knockout partially reversed the growth-inhibitory effects of TBXT loss and modulated a subset of the TBXT-regulated proteome. This included components of a tightly regulated network responsible for interferon signaling upon viral infection or cytokine stimulation. In summary, IGFBP3 may act as a regulator that fine-tunes specific transcriptional effects of TBXT in chordoma cells to create optimal growth conditions.

Our data are consistent with the recent observation that chordoma cells have high ISG scores [80], which reflect the activity of genes directly downstream of the interferon-α/β and γ signaling pathways [84]. Based on comprehensive gene and protein expression analysis, we demonstrated that TBXT inhibition results in markedly lower ISG scores, possibly caused by lowering the transcription factors STAT1 and STAT2, which usually activate ISGs after phosphorylation by JAK family proteins and complex formation. These results have led us to conclude that TBXT directly or indirectly activates the interferon response pathway in chordoma cells and that IGFBP3 fine tunes the TBXT-directed modulation of STAT proteins. When testing drugs that target this pathway on different levels, we found a strong dependence of chordoma cells on JAK2. Since several JAK2 inhibitors are approved, this finding could be immediately applied clinically. This is also supported by the fact that we observed strong interferon signaling in cell lines and chordoma patients and that JAK2 inhibition with fedratinib inhibited the growth of patient-derived chordoma organoids [11]. In addition, targeting STAT3 with experimental compounds has been shown to reduce the viability of chordoma cell lines [85, 86]. Future studies need to determine why chordoma cells exhibit elevated JAK-STAT signaling and to what extent and how TBXT contributes to this characteristic. Another hypothesis could be that genes encoding type I interferons are co-deleted with *CDKN2A* in chordomas [87], causing the cells to increase JAK and STAT levels to compensate for the inability to respond to external cytokines.

In conclusion, we have developed the first selective TBXT protein binders and provide proof of concept that such compounds can target nuclear proteins. In addition, the T-DARPins have allowed us to map the TBXT regulome, which provides a resource that will inform future fundamental and translational studies and has enabled the discovery of an essential signaling axis that can be targeted with clinically approved drugs and thus may represent an immediately actionable therapeutic opportunity in chordoma patients. Finally, the T-DARPins themselves could be developed into clinical drugs for chordoma or other TBXT-expressing cancers if delivery to cells in vivo can be achieved.

## METHODS

### Generation and verification of T-DARPins

The purification of full-length TBXT and the TBXT DBD is described in the Supplementary Methods.

Ribosome display selection: To generate TBXT-specific DARPins, the biotinylated TBXT DBD was immobilized on MyOne T1 streptavidin-coated beads (Pierce) or Sera-Mag neutravidin-coated beads (GE Healthcare) depending on the particular selection round. Ribosome display selections were performed essentially as described [88], yet using a semi-automatic KingFisher Flex MTP96 well platform. The library included N3C-DARPins with the original randomization strategy as reported [32] but contained a stabilized C-cap [89]. Additionally, the library consisted of a mixture of DARPins with randomized and non-randomized N- and C-terminal caps [23, 90]. Successively enriched pools were cloned as intermediates into a ribosome display-specific vector [90]. Selections were performed over four rounds with decreasing concentrations of the biotinylated TBXT DBD and increasing washing steps, and the third round included a competition with the non-biotinylated TBXT DBD to enrich binders with high affinities. The final enriched pool of cDNAs encoding putative DARPin binders was cloned into a bacterial pQE30 derivative vector (Qiagen) via unique *BamHI* and *HindIII* sites as a fusion construct with an N-terminal MRGS(H)_8_-tag and a C-terminal Flag-tag containing a T5 lac promoter and *lacI^q^* for expression control. After the transformation of *E. coli* XL1-blue cells, 380 single DARPin clones selected to bind the TBXT DBD were expressed in 96-well format by adding 1 mM IPTG and lysed by adding B-PER reagent plus lysozyme and nuclease (Pierce). After centrifugation, these crude extracts were used for initial screening to bind the TBXT DBD and full-length TBXT using HTRF and ELISA, which are described in detail in the Supplementary Methods.

#### EMSA

For immobilized metal affinity chromatography (IMAC) purification, 23 identified DARPins were subcloned into a pQIq vector, replacing the MRGS(H)_8_-tag by an MRGS(H)_6_-tag, expressed in small-scale deep-well 96-well plates, lysed with Cell-Lytic B (Sigma), and purified over a 96-well IMAC column (HisPur Cobalt plates, Thermo Scientific), including washing with high-salt (1 M NaCl) and low-salt (20 mM NaCl) phosphate buffer. Elution was performed with PBS containing 400 mM NaCl and 250 mM imidazole. For the EMSA assays, imidazole and NaCl were removed using dialysis devices (3.5K MWCO, Thermo). To increase the amount of the 23 candidate DARPins, large-scale expression and purification were performed. DARPins were expressed on a 200 mL scale. After induction with 1 mM IPTG and incubation for four hours at 37°C, expression cultures were harvested by centrifugation for 10 minutes at 4,000 rpm. Cell pellets were resuspended in sodium phosphate-based Ni-lysis buffer (PBS, 400 mM NaCl, 20 mM imidazole, 10% glycerol, pH 7.4), adding Pierce Universal nuclease, and cells were lysed using sonication. DARPins were purified over a HisTrap FF crude column (GE Healthcare) and desalted over a HiTrap 26/10 desalting column (GE Healthcare) using an ÄKTA Pure L1 system (GE Healthcare). EMSA assays were performed as follows: 72 pmol TBXT DBD were incubated with 135 pmol IMAC-purified and dialyzed DARPins for 30 minutes at room temperature in PBS. To analyze binding of the TBXT DBD to DNA, 1.5 pmol of a double-stranded DNA oligonucleotide containing two palindromic T-box binding elements (5’-CATGAAGGATCCATGAA**<u>TTTCACACCT</u>***<u>AGGTGTGAAA</u>*TTGC-3’; IDT) [10] and 5’ labels with 6-FAM on both strands was added and incubated for 20 minutes at room temperature in the dark. Complex formation was analyzed by native 4% acrylamide gel electrophoresis (Mini-gel, Hoefer) using 0.5 x TBE as running buffer (50 mM Tris hydroxymethyl aminomethane, 50 mM boric acid, 1 mM EDTA, pH 8.0). Migration of the 6-FAM-labeled oligonucleotide was visualized using a fluorescence imager with excitation at 495 nm and emission at 520 nm.

### Dual-luciferase reporter assay

U-CH2, UM-Chor1, and HEK293T + TBXT cells were transduced with 200 µL pLenti6.2-V5/DEST DARPin lentivirus. Medium was changed after 24 hours, and cells were seeded after 72 hours. UCH-2 and UM-Chor1 (8,000 cells/well) and HEK293T + TBXT (3,000 cells/well) were seeded into white, clear bottom 96-well plates, and 24 hours later, cells were transfected with pGL4.34-2X-T-Resp (150 ng/well) and pGL4.73[*hRluc*/SV40] (7.5 ng/well) plasmids using Lipofectamine 3000 (Invitrogen). Non-transfected cells and a single transfection with 150 ng pGL4.34-2X-T-Resp were included as negative controls. After 48 hours, cells were washed once with PBS and lysed using 20 µL Passive Lysis Buffer (Promega) with shaking at 1,000 rpm for 20 minutes at room temperature. Measurements were performed with a Victor X3 plate reader (PerkinElmer) following the Dual-Luciferase Reporter Assay System manual (Promega) instructions. *Firefly* and *Renilla* luciferase signals were corrected by subtracting the respective control samples (non-transfected cells for *Firefly* and *Firefly* cross-talk for *Renilla*). The reporter activity was then calculated by dividing the corrected *Firefly* signal by the corrected *Renilla* signal.

### Cell culture, vectors, and lentiviral transduction

Detailed information on the culturing of cell lines, the generation of cDNAs and lentiviral vectors, and the lentiviral transduction of cell lines is provided in the Supplementary Methods.

### Preparation of protein lysates

For western blotting, IGFBP3 ELISA, and AP-MS, cell pellets were washed once in cold PBS, lysed in RIPA buffer (Thermo Scientific) containing 1 x Halt protease inhibitor cocktail (Thermo Scientific) and 1 x Halt phosphatase inhibitor (Thermo Scientific), and incubated on ice while gently shaking for 30 minutes. Samples were centrifuged at 14,000 g for 15 minutes at 4°C, supernatant was removed, protein concentration was quantified using BCA assay (Pierce), and lysates were stored at −80°C. For DDA-MS, proteins were generated as described above, except cells were harvested by scraping on ice before pelleting. For DIA-MS, proteins were generated as described for DDA-MS with an additional sonication step (BioRupter, Diagenode) for 15 cycles with 30 seconds on/off before submission.

### DARPin IP

For pulldown of Flag-tagged DARPins from lentivirally transduced U-CH2 and UM-Chor1 cells, 5 mg (AP-MS) or 2 mg (western blotting) protein lysate were added to 50 µL Anti-DYKDDDDK (Flag) magnetic agarose beads (prewashed three times, Pierce). After incubation for one hour at 4°C on a rotating shaker, the supernatant was discarded, and beads were washed three times for five minutes with PBS containing 0.05% Tween-20. Elution was performed with 80 µL 3 x Flag peptide (1.5 mg/mL in PBS, Sigma-Aldrich) and by incubating for 20 minutes at room temperature in a ThermoMix Comfort shaker (1,400 rpm, Eppendorf). The supernatant was separated from the magnetic beads, and 30 µL eluate or input lysate were used for western blotting, or the entire eluate was submitted to the DKFZ Mass Spectrometry-Based Protein Analysis Core Facility for AP-MS analysis.

For pulldowns from HEK293T cells, these were co-transfected with pShuttle CMV-IRES-GFP encoding a DARPin and pLEX307 containing no cDNA, TBXT-HA, TBXT-R16L-HA, TBXT-H171R-HA, TBXT-G177D-HA, or TBXT-ΔDBD-HA. Cells were harvested by scraping, lysed in IP Lysis Buffer (25 mM Tris-HCl, 150 mM NaCl, 1 mM EDTA, 1% Triton, 5% glycerol, pH 7.4) containing 1 x Halt protease and phosphatase inhibitors, and centrifuged at 13,000 x g for 10 minutes. Protein lysate (1 mg) was added to 30 µL Anti-DYKDDDDK (Flag) magnetic agarose beads (prewashed three times) and incubated overnight at 4°C on a rotating shaker. The supernatant was discarded, and beads were washed four times for five minutes with PBS + 0.05% Tween-20. Elution was performed with 20 µL 3 x Flag peptide (1.5 mg/mL in PBS, Genaxxon BioScience) and by incubating for 20 minutes at room temperature in a Thermomix shaker (1,400 rpm, Eppendorf). The supernatant was separated from the magnetic beads, and 10 µL eluate or 50 µg input lysate were used for western blotting.

### Western blotting

Western blotting was performed using 10 µg protein lysate (25 µg for detecting DARPins) mixed with 4 x Laemmli sample buffer (Bio-Rad) and diluted to 1 x with dH_2_O and addition of β-mercaptoethanol (Sigma-Aldrich) to a final concentration of 10%. The samples were heated to 95°C for 10 minutes, cooled, and loaded on a 4–20% Mini-PROTEAN TGX gel (Bio-Rad). The gels were run for 10 minutes at 90 V and 45 minutes at 150 V. Proteins were transferred to PVDF membranes (Bio-Rad) using the Trans-Blot Turbo Transfer System (Bio-Rad) for 30 minutes. Membranes were blocked for one hour at room temperature with 5% milk in TBST (1 x tris-buffered saline, 0.1% Tween-20), incubated with primary antibodies overnight at 4°C with gentle rocking, and washed three times for 10 minutes each in TBST before incubation with a secondary antibody for one hour at room temperature. After three additional washing steps in 1 x TBST, blots were imaged using the Odyssey CLx (Li-Cor) or the ChemiDoc (Bio-Rad) using SuperSignal West Atto Ultimate Sensitivity Substrate (A3855, Thermo Scientific). To visualize IGFBP3, membranes were blocked with Intercept TBS buffer (Li-Cor). Primary and secondary antibodies are provided in **Supplementary Table 5**.

### Mass spectrometry sample preparation, raw data acquisition, and data preprocessing

Information on AP-MS, DDA-MS, and DIA-MS sample preparation, raw data acquisition, and data preprocessing is provided in the Supplementary Methods.

### Mass spectrometry data analysis

#### DDA-MS

LFQ matrices calculated by MaxQuant were imported into R using the DEP package [91]. Non-targeting control samples (E3_5 DARPin) were used as baselines. First, protein groups were filtered to analyze only those detected at least once for all three replicates in each group (E3_5 and T-DARPin). Second, the protein groups with a median LFQ below the 0.05 quantile for all LFQ values (control or treatment) were removed. Finally, the filtered LFQ matrix was normalized using variance stabilizing normalization [92]. Missing values were imputed using the MinDet method as implemented in the imputeLCMD::impute.MinDet function. The minimum was estimated by using the 0.01 quantile.

DIA-MS: The DIA-MS data were processed similarly to the DDA-MS data, except that only proteins detected at least twice in each group were kept, and the LFQ quantile cutoff was reduced to 0.015.

#### Differential protein expression

Protein levels were compared by fitting linear models using the *limma* package in R and a robust empirical Bayes estimation with a variance trend [93]. The following linear model was used, where *DARPin* represents a categorical variable encoding the E3_5 or T-DARPin conditions:

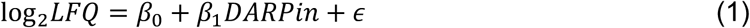

Differential expression and its statistical significance were calculated by a moderated t-test. For each comparison of a T-DARPin with the E3_5 condition, proteins with at least 1.5-fold up- or downregulation and an FDR of less than 5% were selected as significantly deregulated.

#### AP-MS

After data preprocessing, the proteins detected in E3_5 pulldowns in both cell lines were filtered out to focus specifically on the proteins detected in T-DARPin pulldowns. Second, the same linear model (equation 1), moderated t-test, and FDR filter were applied to these reduced datasets as for the differential protein expression analysis, but no fold-change filter was applied. Finally, the fold change for each protein in a sample (T-DARPin treatment in a given cell line) was normalized to the highest fold change in the same sample to calculate the normalized enrichment.

Multifactorial linear regression with interactions: DIA-MS data were analyzed separately for the effect of TBXT inhibition in the context of IGFBP3 knockout. First, we merged the datasets for co-detected proteins in the sgNTC and sgIGFBP3 conditions. Second, we applied the following model, where *IGFBP3* and *DARPin* are categorical variables encoding the sgNTC/sgIGFBP3 and NTC/T-DARPin D4 treatments, respectively, and extracted the effect size and p-value from the moderated t-test for the interaction term alone:

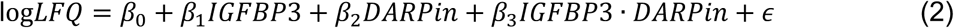

#### STRING network

Functional association networks of the proteins enriched in the AP-MS analysis were analyzed with the STRING database version 12.0 [94] using the following settings: network type selected as full STRING network, network edges showing evidence, all active interaction sources selected, interaction score set at medium confidence (0.4), and disconnected nodes hidden.

### RNA preparation

Cell pellets were resuspended with 350 μL RLT buffer on ice, lysed with a 26-gauge needle, and processed with the RNAeasy Mini Kit (Qiagen), including a DNAse I digestion step. Eluted RNA was immediately frozen at −80°C with 1 µL RNaseOUT (Invitrogen) after quantification with a NanoDrop One system (Thermo Scientific).

### Quantitative RT-PCR

For each condition, 1 µg RNA was reverse-transcribed using the High-Capacity cDNA Reverse Transcription Kit (Applied Biosystems). cDNA was diluted 1:20 and combined with primers for the target genes and SYBR Green master mix (Bio-Rad). Amplification was performed using standard conditions on a CFX96 Real-Time system (Bio-Rad). Relative mRNA expression was quantified using the 2^-ΔΔCT^ method [95] after obtaining CT values from the CFX Manager software (Bio-Rad). Primer sequences are provided in **Supplementary Table 6**.

### RNA-seq and differential gene expression analysis

UM-Chor1 cells were transduced with pLenti6.2-V5/DEST encoding E3_5 or T-DARPins A2, B1, and D4, each in triplicate, selected with blasticidin (4 µg/mL) for 10 days, and cultured for two days without selection before RNA isolation. RNA was quantified and checked for purity on a TapeStation (Agilent). High-quality RNA (RIN > 8.0) was submitted to the DKFZ Next-Generation Sequencing Core Facility, where sequencing librariers were prepared using Illumina TruSeq Stranded Kit and sequenced on an Illumina HiSeq 4000 platform at 100 cycles (paired end). Details on RNA-seq and data preprocessing are provided in the Supplementary Methods.

Differential gene expression: Differential expression analysis was performed using the DESeq2 package [96]. Gene-specific read count data were imported into an R environment. Active genes were selected with the zFPKM method using the suggested zFPKM cutoff of −3 [96]. Biological replicates were analyzed for batch effects using the sva::ComBat function and principal component analysis. The batch term *Replicate* was introduced into the model to adjust for batch effects:

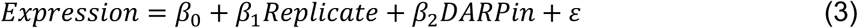

Next, differential expression analysis was performed using E3_5 samples as the baseline for all comparisons. The FDR was set to 1% using independent hypothesis weighting [97]. Misleading fold changes of genes with low expression or high dispersion were adjusted using adaptive empirical Bayes shrinkage [98]. For each comparison, genes with at least 1.5-fold up- or downregulation with an FDR less than 1% were selected as significantly regulated.

### GSEA

#### T-DARPin experiments

Fast pre-ranked GSEA was performed with the *fgsea* package in R [99] using the human MSigDB Hallmark, GO_BP version 2023.2.Hs, and Enrichr ARCHS^4^ Tissue gene collections [100, 101]. An FDR-adjusted p-value of 0.05 was used to infer significantly enriched gene sets. Due to the large number of significantly deregulated gene sets in the GO_BP and ARCHS^4^ Tissue collections, enrichment levels were simplified for representation [102]. GO_BP terms were clustered based on semantic similarity. ARCHS^4^ Tissue gene sets were clustered based on normalized enrichment scores, but gene set identifiers were simplified in word clouds. Word sizes were determined by the product of the absolute normalized enrichment score and the negative log_10_ of the p-value.

Patient samples: Gene expression levels normalized as transcripts per million (TPM) for 1,077 samples from 1,008 patients representing 233 cancer types [60] were subjected to gene set variation analysis [103] using Hallmark gene sets. The minimum size of a resulting gene set after gene identifier mapping to expression data was set to 5 to improve the reliability of the enrichment scores, calculated as the difference between the maximum and minimum deviations of the random walk from the origin using a modified Kuiper statistic. The resulting scores were separated into chordoma (12 samples) and others (1,065 samples). Mean values and 95% confidence intervals were calculated for each Hallmark gene set for each group.

### Integrative analysis of transcriptome and proteome data

#### Definition of commonly deregulated genes and proteins

At the transcript level, commonly deregulated genes were defined by a significant change in the same direction by all three T-DARPin treatments. At the protein level, more permissive criteria were used. We aimed to identify proteins significantly altered in the same direction by at least two T-DARPin treatments, eliminating discordant cases deregulated in one direction by two treatments but in the opposite direction by the third treatment. We defined two mathematical functions to apply this logic. First, we discretized the direction of deregulation into three levels (−1, 0, and 1) for each protein group *g* in each T-DARPin treatment *t* (equation 4), where sgn is the signum function, and *f* and *p* represent the log_2_(fold change) and FDR, respectively:

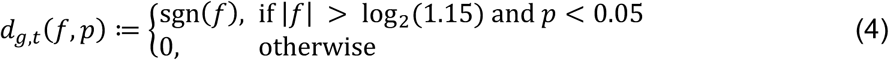

Second, we encoded the “commonly deregulated” property as a Boolean value for each protein group using a discretized deregulation function (equation 5), where T is the set of all T-DARPin treatments:

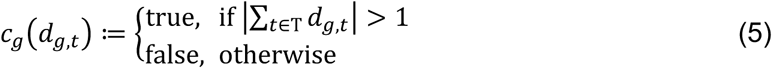

Data harmonization: DEPs and DEGs were matched by mapping UniProtKB protein identifiers to Ensembl gene identifiers. GENCODE 19 gene names were updated to the GENCODE 45 annotation, originally created for GRCh38 reference chromosomes and mapped back to the GRCh37 primary assembly.

Drug targetability: Potentially druggable human genes were previously identified by Jiang et al. [44]. We used Table S9 from this publication to annotate our integrated transcriptome and proteome data with target development level and gene family information.

### Spheroid formation assays

The 3D culture of chordoma cells in Matrigel was performed as follows: eight-well glass slides were coated with 60 µL Matrigel (Corning, Basement Membrane Matrix, LDEV-Free) per well, and 2,000 chordoma cells per well were seeded in complete chordoma medium containing 4% Matrigel. The medium was changed once a week, and after 34 (U-CH2) or 60 (U-CH12) days, spheroids bigger than 50 µm were counted with a bright field microscope.

For single spheroid growth assays, 5,000 cells per well of UM-Chor1-iDARPin, UM-Chor1-EV, UM-Chor1-IGFBP3, U-CH2-EV, U-CH2-IGFBP3, or UM-Chor1-Cas9 transduced with sgNTC, sgTBXT, or sgIGFBP3 were seeded in an ultra-low attachment 96-well plate (Corning) and allowed to aggregate into one spheroid per well for seven to 10 days. Half-medium exchanges were performed one to two times a week. DARPin, IGFBP3, or empty vector (EV) expression was induced using 0.5 µg/mL doxycycline or sterile dH_2_O as a negative control. Spheroids were imaged every seven days using a Lionheart FX automated microscope (BioTek) in 96-well acquisition format. Guiding beacons were set for each well, and spheroid images were acquired at four-fold magnification using bright field illumination (LED 7, integration time 220, gain 1). A 10-step (250 µm per step) z-stack was acquired for each spheroid and compressed into a projection image (z-proj). Z-proj images were imported into FIJI version 1.53C, and the spheroid area was calculated with a customized FIJI macro and reported as µm^2^ x 10^5^. To measure their viability, spheroids were transferred using wide-bore pipette tips to white opaque 96-well plates (Corning), and CellTiter-Glo 3D reagent assay (Promega) was used following the manufacturer’s instructions. Luminescence was acquired with an Envision Multimode plate reader (PerkinElmer), and raw units (x 10^7^) or raw units normalized to EV controls were reported.

### Cell viability, cell cycle, and apoptosis analysis

The quantification of viable cells by trypan blue staining and MTS assay and the analysis of cell cycle and apoptosis by flow cytometry are described in the Supplementary Methods.

### Mouse experiments

NOD.Cg-*Prkdc^scid^ Il2rg^tm1Wjl^*/SzJ (NSG) mice were housed in the DKFZ Center for Preclinical Research. All animal procedures were approved by the regional authority in Karlsruhe, Germany (reference number 35-9185.81/G-51/21) and performed according to federal and institutional guidelines. U-CH1 cells (2 x 10^5^ per well) were seeded in six-well plates, spin-infected with 10 uL concentrated pLenti6.2 E3_5 or D4 lentivirus per well after three days, selected with 6 µg/mL blasticidin for one week, and allowed to recover in selection-free media for four days (**Supplementary Fig. 3g**). U-CH1 cells expressing E3_5 or D4 were harvested, washed, and resuspended in DPBS. 1 x 10^6^ cells in 55 µL DPBS were mixed with 55 µL Matrigel (Corning, Growth Factor Reduced Basement Membrane Matrix, LDEV-Free), of which 100 µL were injected subcutaneously into the flanks of isoflurane-anesthetized, female, six to eight-week-old NSG mice provided by the DKFZ Center for Preclinical Research. Each cell line was applied to both flanks of three mice, resulting in six E3_5 tumors and six D4 tumors. The animals were monitored closely, and the width and length of the tumors were measured with a caliper. The tumor volume was calculated using the formula (length x width x width)/2. The mice were sacrificed after reaching the permitted criteria of a maximum tumor size of 2 cm in one diameter and a maximum tumor volume of 1.5 cm^3^, and tumors were processed for histopathologic, protein, and RNA analyses.

### Immunofluorescence

Immunofluorescence analyses of UM-Chor1 cells and spheroids are described in the Supplementary Methods.

### IGFBP3 protein analyses

Information on the quantification of IGFBP3 using ELISA and the analysis of IGFBP3 glycosylation and secretion is provided in the Supplementary Methods.

### Drug treatment of cell lines

Fedratinib (#HY-10409, MedChemExpress), AZD1480 (#SML1505, Sigma), fludarabine (#F9813, Sigma), filgotinib (#HY-18300, MedChemExpress), gilteritinib (#S7754, Selleckchem), pacritinib (#S8057, Selleckchem), and afatinib (#SML3073, Sigma) were dissolved to 10 mM in sterile DMSO (Sigma) and stored at −80°C. Before use, compounds were redissolved for 10 minutes at 37°C, vortexed, and inspected for the absence of precipitates. Compounds were never subjected to multiple freeze-thaws. 1,000 cells per well were plated in technical and biological triplicate into collagen-coated white clear-bottom 96-well plates and allowed to adhere for 24 hours before a seven-day drug treatment. For the initial drug screen, compounds were serially diluted from 10 µM to 1 nM. A T_0_ plate for calculating the GR was measured using CellTiter-Glo (Promega) following the manufacturer’s instructions. After seven days, plates were measured using CellTiter-Glo (Promega) following the manufacturer’s instructions. Luminescence was acquired using an Envision Multimode plate reader (PerkinElmer). GR values were obtained with grcalculator.org [69] and graphed using GraphPad Prism. Colony formation assays (14 days, 1 µM drug) were performed as described previously [104] using normal growth medium and an initial seeding density of 10,000 cells per 6-well for U2OS, and 25,000 cells per 6-well for U-CH1, UCH2, UM-Chor1, and JHC7 cells. For apoptosis measurements, cells were treated with 3 µM drug for 4 or 7 days. Information on the flow cytometry based assays is provided in the Supplementary Methods.

### Statistical analysis

Statistical analyses were performed with GraphPad Prism version 10.2.2 or R version 4.3.2. For the analysis of experiments, averages and standard errors of the mean were generated from biological replicates, and statistical significance was determined using one-way ANOVA and the post-hoc Dunnett’s test for multiple comparisons or unpaired t-test.

## Data availability

Raw and preprocessed RNA-seq data from cell lines have been deposited in the Gene Expression Omnibus under the accession number GSE276715 ((https://www.ncbi.nlm.nih.gov/geo/query/acc.cgi?acc=GSE276715). Sequencing data from Horak et al. [60] are available with restricted access under accession number EGAS00001004813 [https://www.ebi.ac.uk/ega/studies/EGAS00001004813]. The mass spectrometry data have been deposited in the ProteomeXchange Consortium via the PRIDE partner repository with the dataset identifier PXD055982.

## ACKNOWLEDGEMENTS

We thank Anna Imhof and Jana Kress; the DKFZ Light Microscopy, Mass Spectrometry-Based Protein Analysis, Next-Generation Sequencing, Omics IT and Data Management, and Tumor Models Core Facilities; and the European Molecular Biology Laboratory Protein Expression and Purification Core Facility for technical support. We thank Sebastian Dieter, Michael Wegert, Leila Martins, Usman Shah Gilani, Michelle Michelhans, and the Chordoma Foundation for insightful discussions. This study was supported by grant 70113002 from the German Cancer Aid and grant 2019.043.01 from the Wilhelm Sander Foundation.

## AUTHOR CONTRIBUTIONS

C.S.U., M.G., S.F., and C.S. conceptualized the study; J.M., B.D., J.V.S., and A.P. performed the T-DARPin selection and validation; C.E. performed bioinformatic analyses of RNA-seq and proteome data; C.S.U., M.G., K.-S.L., F.I., P.S., S.D., A.K., P.W., and J.H. performed functional experiments with cell lines; A.P. provided essential methodologies; T.F.E.B and K.M. provided essential resources. C.S.U. and C.S. wrote the original draft with major input from C.E., A.P., and S.F. All co-authors reviewed and approved the final version of the manuscript.

## Notes

**Conflict-of-interest disclosure:** A.P. is a co-founder and shareholder of Molecular Partners AG, which is commercializing the DARPin technology. S.F. has had a consulting or advisory role and received honoraria, research funding, and/or travel/accommodation expenses funding from the following for-profit companies: Amgen, AstraZeneca, Bayer, Eli Lilly, Pfizer, PharmaMar, and Roche. C.S.U., M.G., J.M., B.D., J.V.S., A.P., S.F., and C.S. have filed a patent (EP 23184301.2) based on the results described in this manuscript. The other authors declare no potential conflicts of interest.

### Competing Interest Statement

A.P. is a co-founder and shareholder of Molecular Partners AG, which is commercializing the DARPin technology. S.F. has had a consulting or advisory role and received honoraria, research funding, and/or travel/accommodation expenses funding from the following for-profit companies: Amgen, AstraZeneca, Bayer, Eli Lilly, Pfizer, PharmaMar, and Roche. C.S.U., M.G., J.M., B.D., J.V.S., A.P., S.F., and C.S. have filed a patent (EP 23184301.2) based on the results described in this manuscript. The other authors declare no potential conflicts of interest.

