## Supplementary Information for "Selective targeting of TBXT with DARPins identifies regulatory networks and therapeutic vulnerabilities in chordoma"

**Supplementary Figure 1**

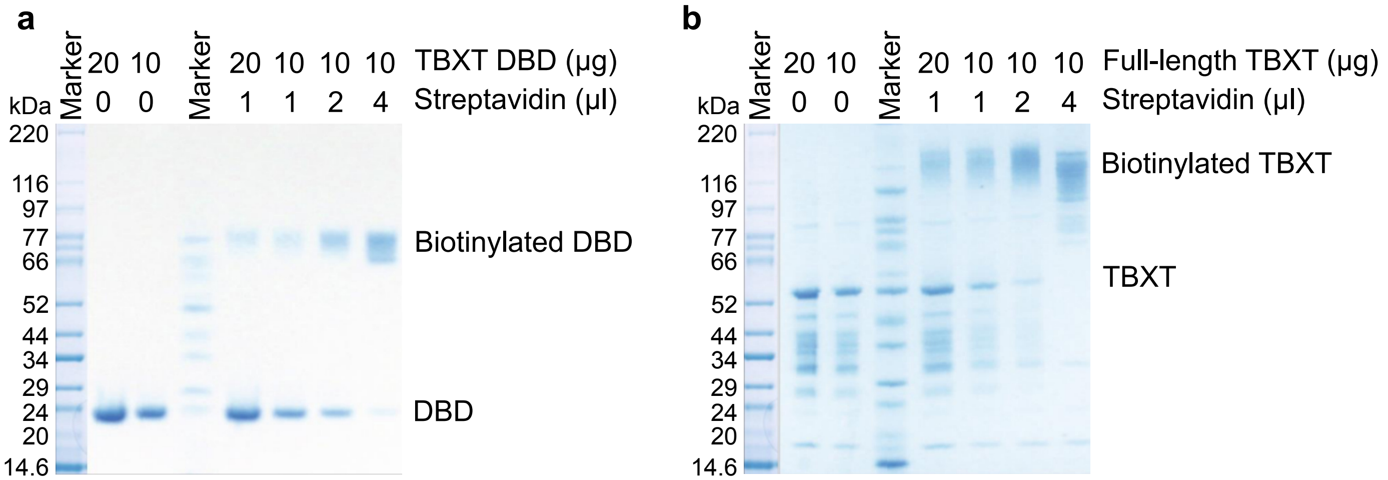


**Supplementary Figure 1. Generation of TBXT-binding DARPins. (a, b)** Coomassie blue-stained SDS-PAGE gels demonstrating biotinylation of the purified recombinant TBXT DBD (a) and full-length TBXT (b) used for the selection of TBXT-binding DARPins. The TBXT DBD was used as the target protein for ribosome display, and full-length TBXT was used in addition to the DBD for the following selection and verification experiments.

**Supplementary Figure 2**

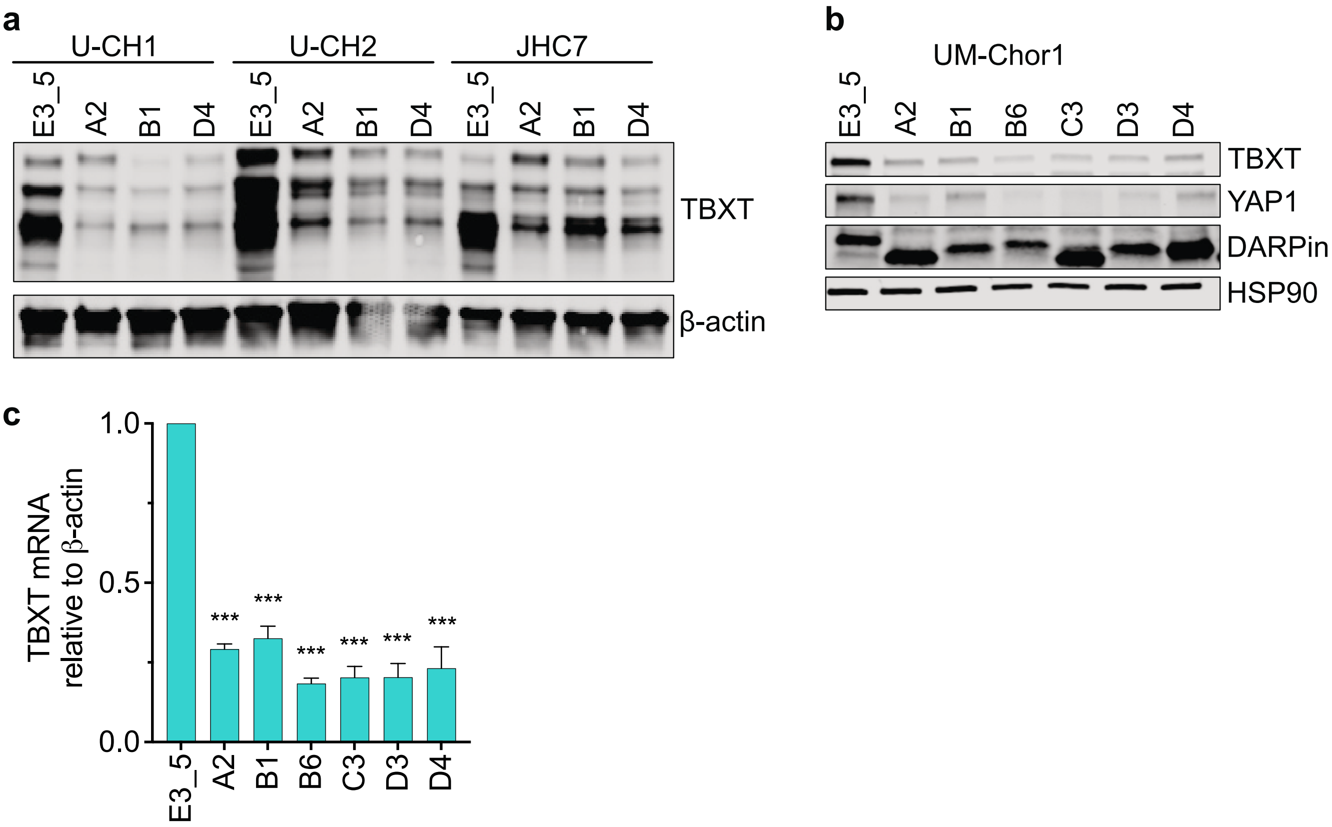


**Supplementary Figure 2. Specificity and activity of T-DARPins. (a)** Western blot of U-CH1, U-CH2, and JHC7 chordoma cell lines 12 days after transduction with the three indicated T-DARPins or E3_5. One representative blot of three independent experiments is shown. **(b)** Western blot of UM-Chor1 cells 12 days after transduction with the six lead T-DARPins or E3_5 with antibodies detecting TBXT and the TBXT downstream effector YAP1. One representative blot of three independent experiments is shown. **(c)** *TBXT* mRNA expression measured by quantitative RT-PCR in UM-Chor1 cells stably expressing the six lead T-DARPins or E3_5. One-way ANOVA with Dunnett's test for multiple comparisons; mean ± SEM of three biological replicates. *p ≤ 0.05, **p ≤ 0.01, ***p ≤ 0.001.

**Supplementary Figure 3**

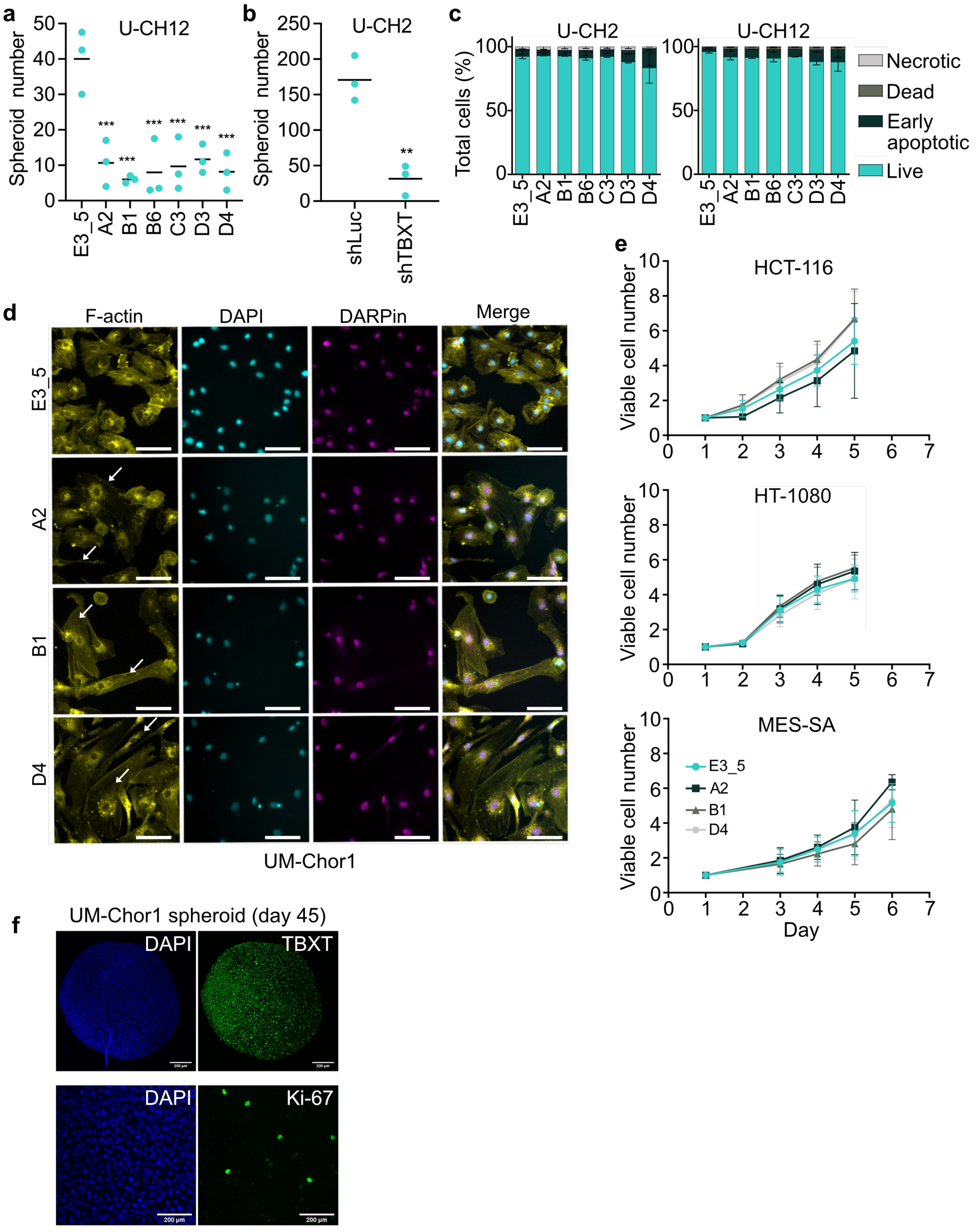

**Supplementary Figure 3 continued**

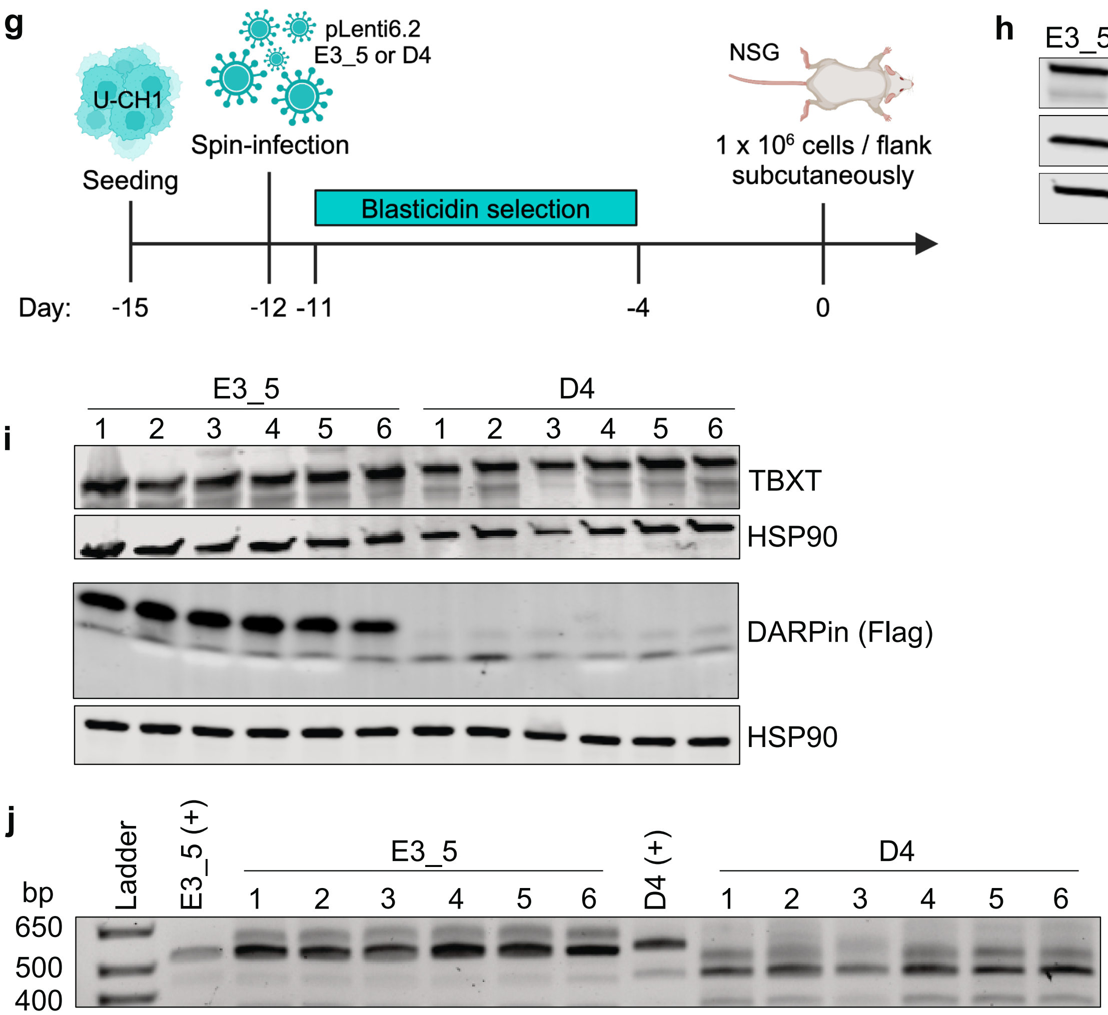


**Supplementary Figure 3. Cellular effects of T-DARPin expression. (a)** 3D Matrigel cultures of U-CH12 cells. Spheroids were counted on day 60 after lentiviral transduction with the indicated DARPins. One-way ANOVA with Dunnett's test for multiple comparisons; mean of three biological replicates. *$p \le 0.05$, **$p \le 0.01$, ***$p \le 0.001$. **(b)** Same experimental setup as in (a) but with lentiviral transduction of an shRNA targeting TBXT or shLuc. Unpaired t-test; **$p \le 0.01$. **(c)** Apoptosis measured by flow cytometry after staining with annexin V and PI 10 days after lentiviral transduction of U-CH2 and U-CH12 cells with the indicated DARPins. Mean ± SEM of two biological replicates. Necrotic, $PI^+$/annexin$V^-$; early apoptotic, $PI^-$/annexin $V^+$; dead, $PI^+$/annexin $V^+$; live, $PI^-$/annexin $V^-$. **(d)** Immunofluorescence of UM-Chor1 cells 10 days after transduction with DARPins A2, B1, D4, or E3_5. Cells were stained with F-actin-specific phalloidin (yellow), nucleus-specific DAPI (teal), and DARPin-specific anti-His (magenta) antibodies. The original images were pseudo-colored for better visualization. White arrows depict pancake-like and spindle-shaped cells. Scale bar, 100 μm. **(e)** Number of viable cells measured by MTS assay relative to day 1 over time of the TBXT-negative cell lines HCT116 (colon cancer), HT-1080 (fibrosarcoma), and MES-SA (uterine sarcoma) expressing DARPins E3_5, A2, B1, or D4. **(f)** Confocal microscopy images of UM-Chor1 spheroids grown for 45 days in ultra-low attachment plates and stained with anti-TBXT and anti-Ki-67 antibodies and DAPI. **(g)** Schematic of the timeline of the preparation of U-CH1 cells before injection into NSG mice. Created in BioRender. Fröhling, S. (2024) BioRender.com/c53k096. **(h)** Western blot of U-CH1 cells transduced with E3_5 and T-DARPin D4 immediately before injection into NSG mice. **(i)** Western blot of lysates from 12 U-CH1 tumors (six transduced with E3_5 and six transduced with T-DARPin D4) at the endpoint of the xenotransplantation experiment. **(j)** PCR from mRNA of 12 U-CH1 tumors (six transduced with E3_5 and six transduced with T-DARPin D4) at the endpoint of the xenotransplantation experiment with primers for amplifying the DARPin mRNAs. The positive controls E3_5 (+) and D4 (+) were amplified from DARPin mRNA extracted from transduced U-CH1 cells immediately before implantation.

**Supplementary Figure 4**

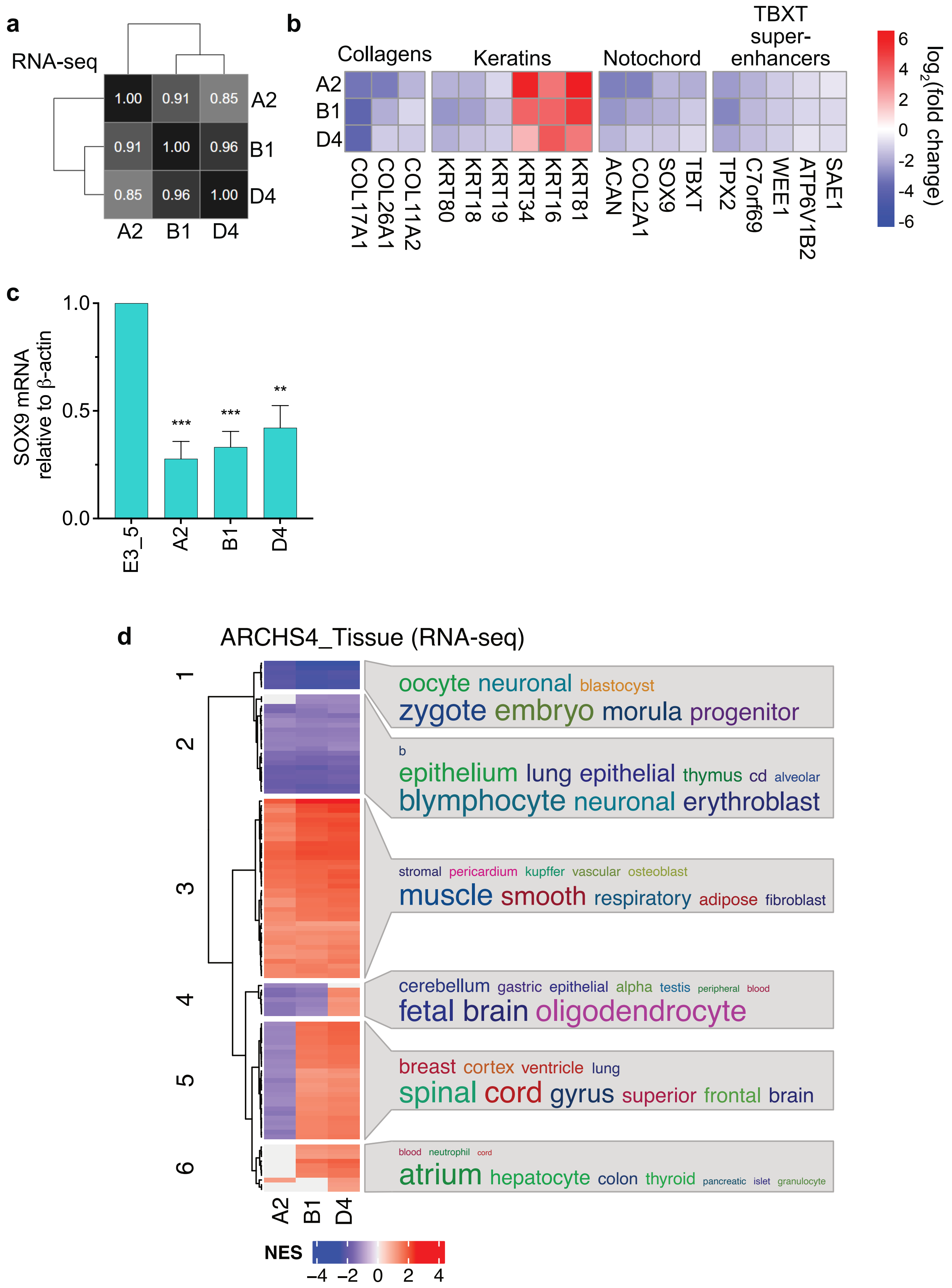

**Supplementary Figure 4 continued**

**e**

GO_Biological Process (RNA-seq, downregulated processes)

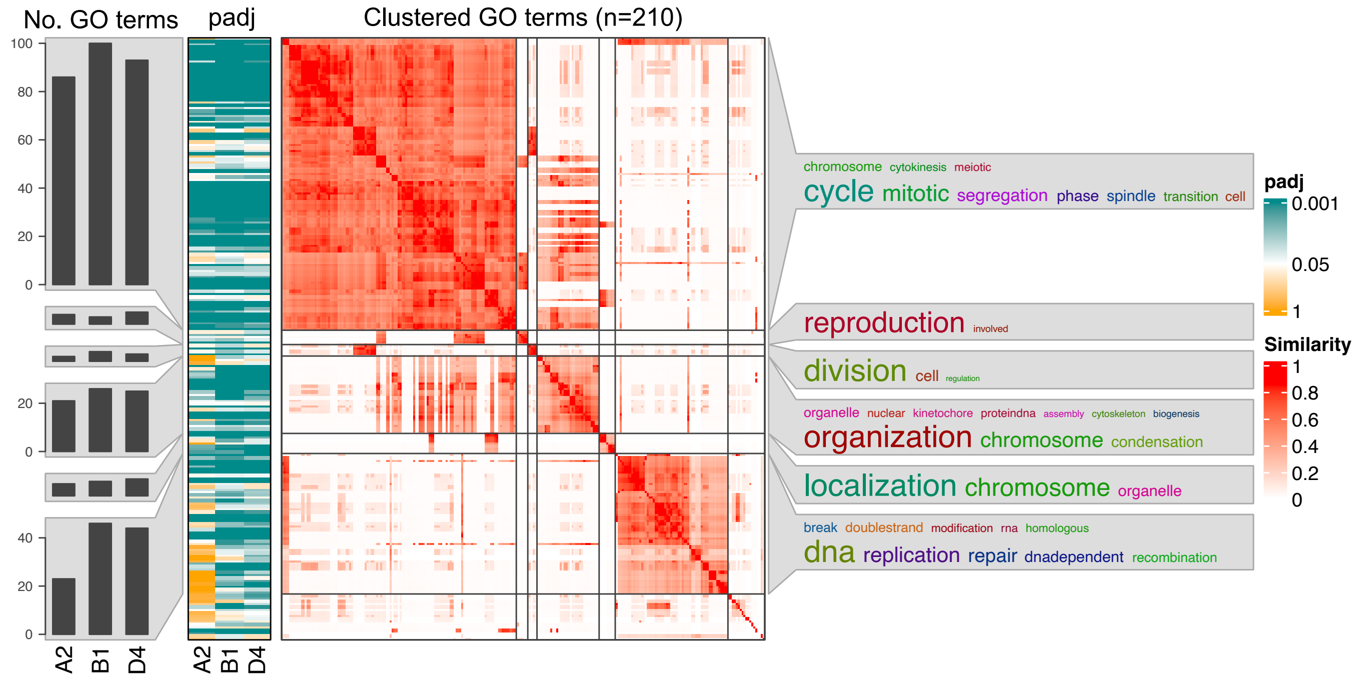


**f**

GO_Biological Process (RNA-seq, upregulated processes)

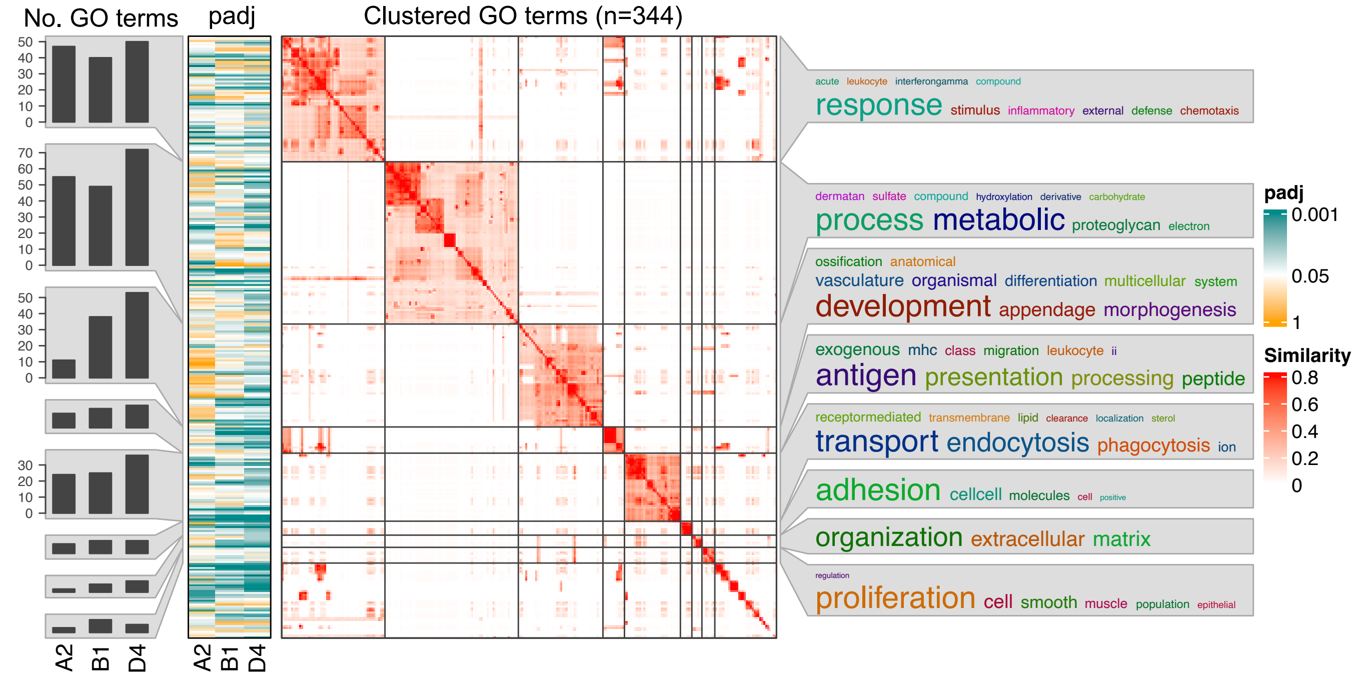

**Supplementary Figure 4 continued**

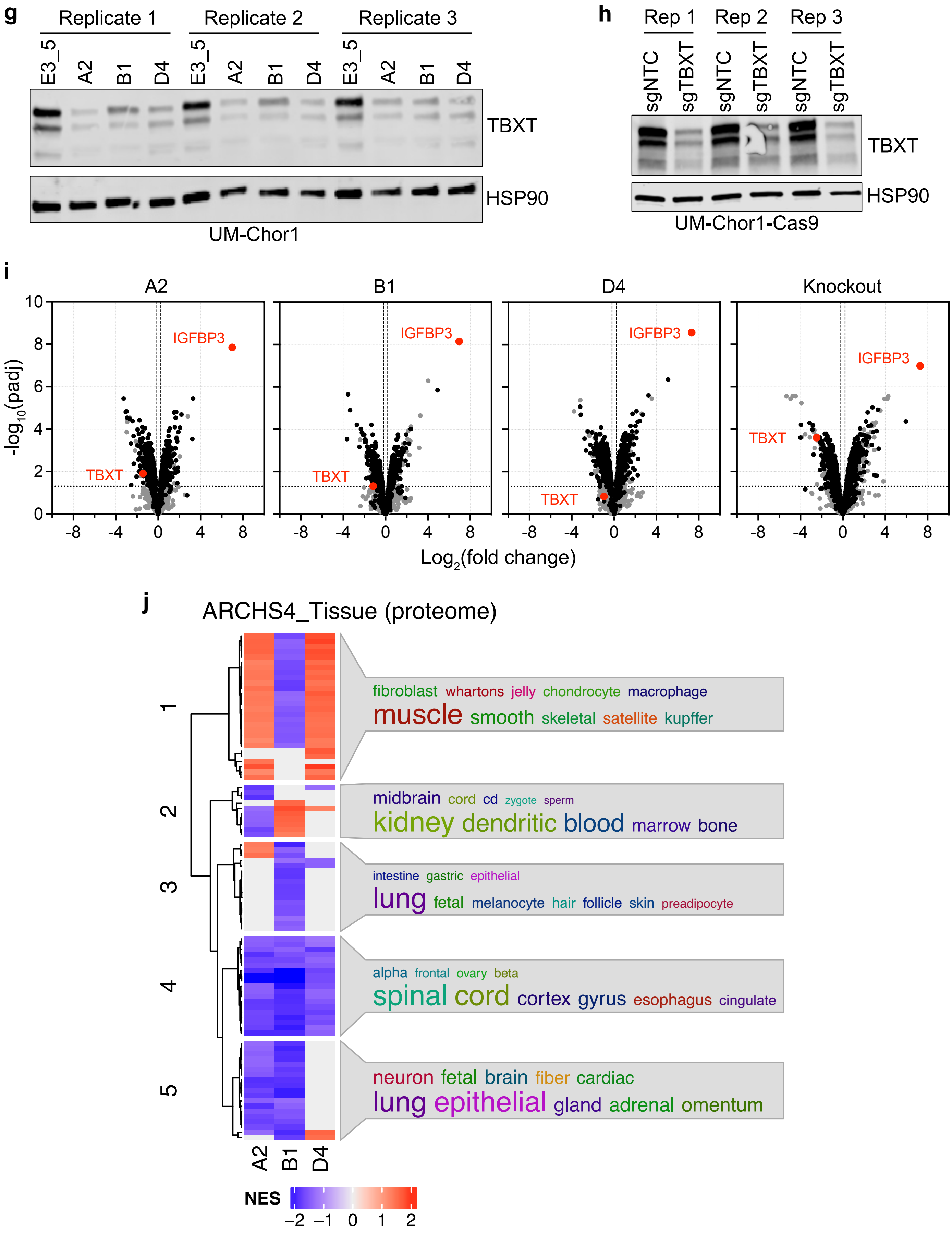

**Supplementary Figure 4 continued**

k

GO_Biological Process (proteome, downregulated processes)

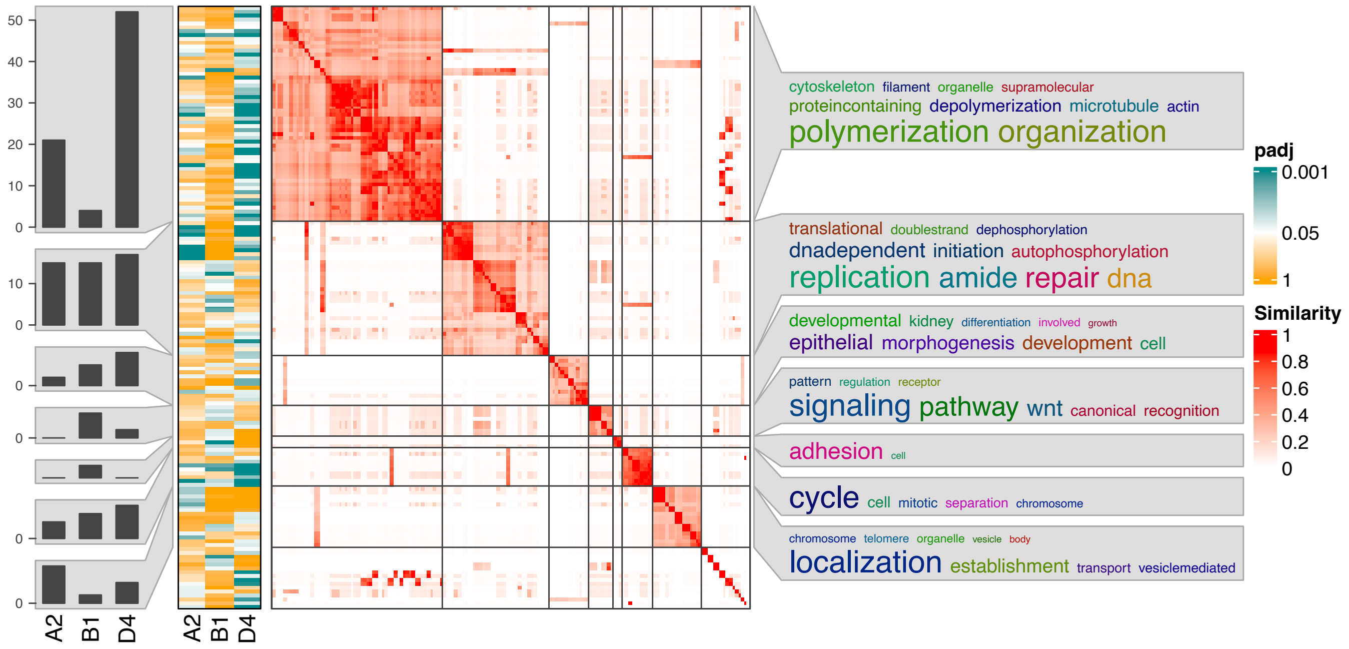


l

GO_Biological Process (proteome, upregulated processes)

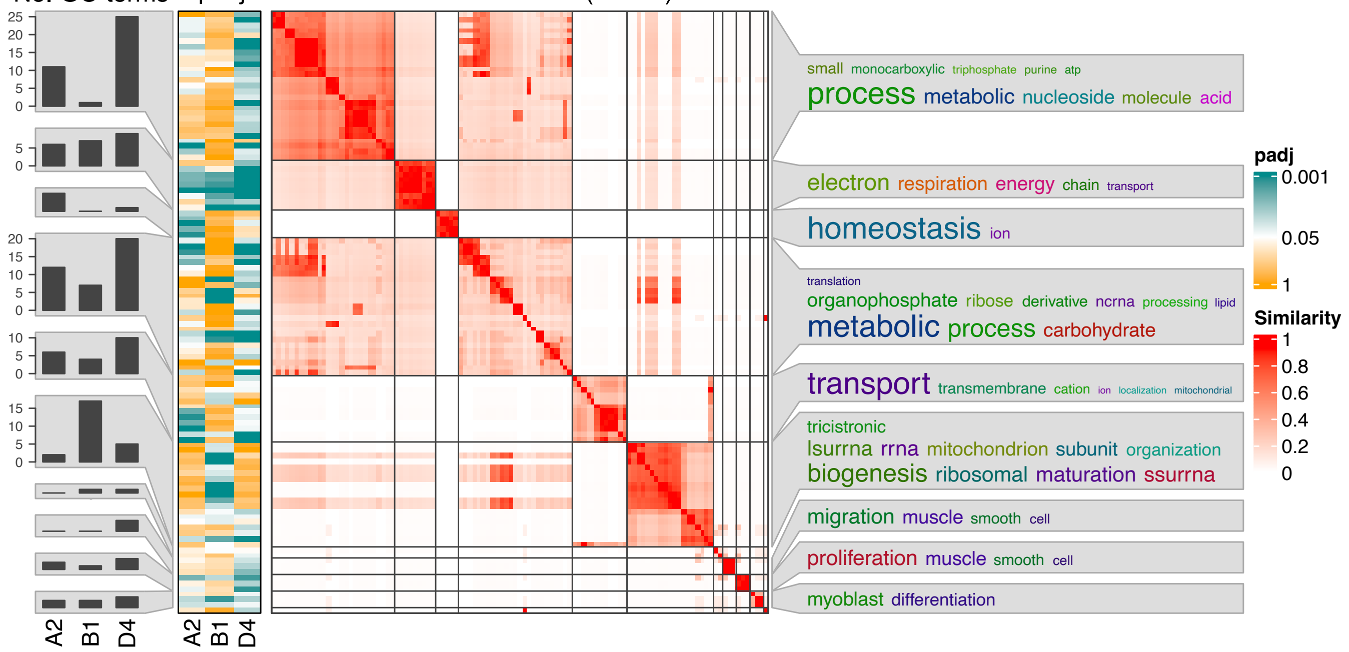

**Supplementary Figure 4 continued**

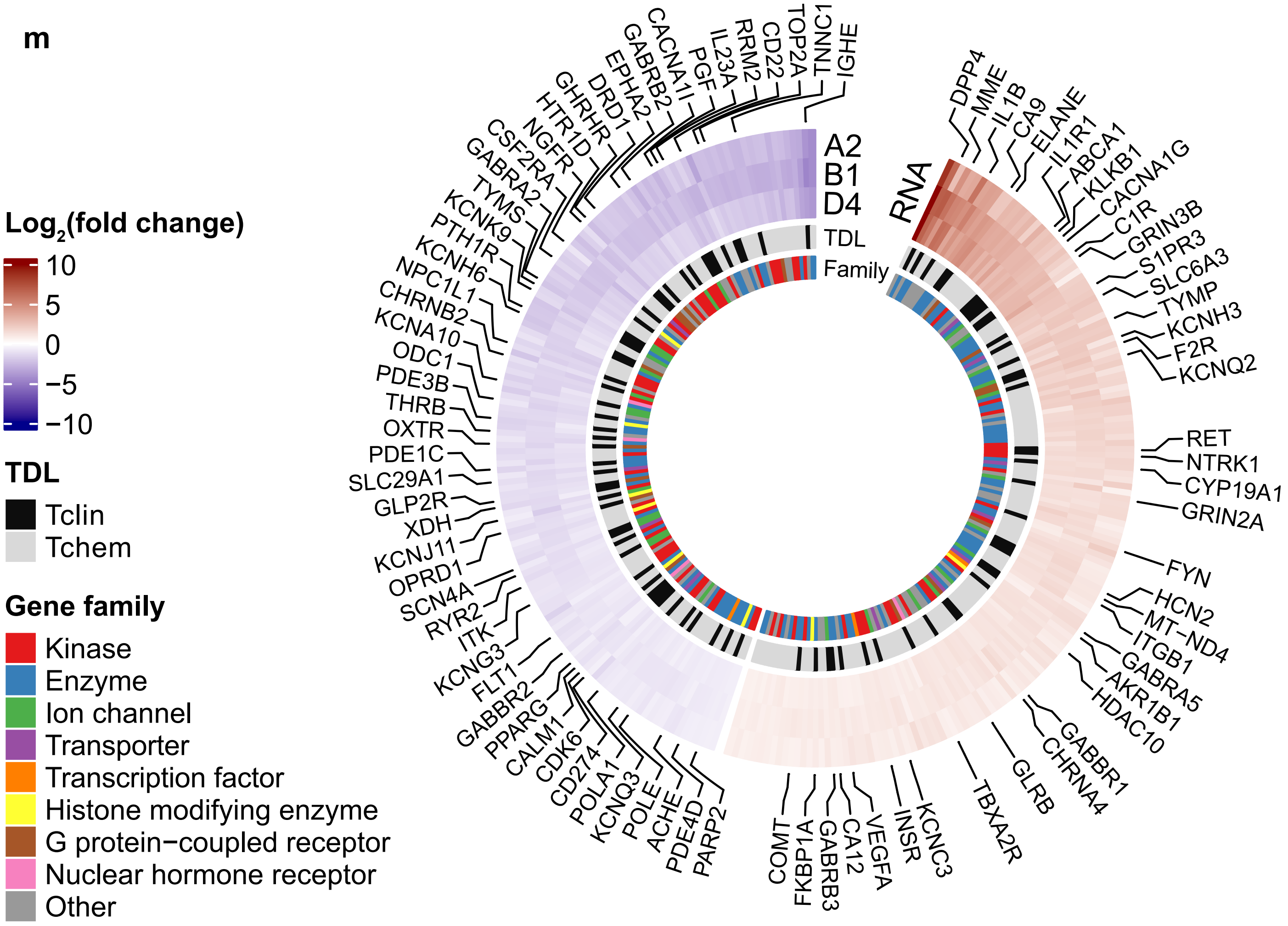


**Supplementary Figure 4. TBXT regulome determined by T-DARPins in UM-Chor1 cells. (a)** Pearson correlation coefficient matrix for pairwise similarity of differential expression levels (log2 (fold change)) in the context of three T-DARPin conditions. RNA-seq of UM-Chor1 cells 12 days after transduction with T-DARPins A2, B1, and D4 and E3_5 performed in biological triplicates. Differential expression was calculated using the DESeq2 workflow comparing gene expression in the context of each T-DARPin treatment to E3_5. **(b)** Heatmap of RNA-seq data depicting genes known to be regulated by TBXT or chordoma-relevant genes. **(c)** *SOX9* mRNA expression measured by quantitative RT-PCR in UM-Chor1 cells stably expressing the three T-DARPins or E3_5. One-way ANOVA with Dunnett's test for multiple comparisons; mean + SEM of three biological replicates. *p ≤ 0.05, **p ≤ 0.01, ***p ≤ 0.001. **(d)** Fast preranked GSEA of RNA-seq data with the ARCHS$^4$ Tissue gene collection. Gene set names were separated into words and represented in word clouds based on the frequency of occurrence of each word. Common words such as "cell" or "tissue" were omitted. NES, normalized enrichment score. **(e)** Fast preranked GSEA of RNA-seq data with the GO_BP gene collection focused on downregulated pathways. The left panel shows the number of significant GO terms (adjusted p-value [padj] < 0.05) in each cluster for each T-DARPin (column); the middle panel indicates the padj as a heatmap for each GO term (row) and T-DARPin (column); and the heatmap in the right panel shows the semantic similarity clustering of GO terms by '"binary_cut". The number of GO terms clustered (n = 210) is shown above this heatmap. The enriched GO terms are summarized per cluster as word cloud by frequency of occurrence as in panel (d). **(f)** Fast preranked GSEA of RNA-seq data with the GO_BP gene collection focused on upregulated pathways. Please see panel (e) for a description. **(g, h)** Western blot of UM-Chor1 cell lysates prepared in triplicate for DDA-MS**.** Cells were transduced 14 days earlier with T-DARPins A2, B1, or D4 or E3_5 (g) or with sgNTC or sgTBXT (h). **(i)** Volcano plots of proteins detected by DDA-MS in UM-Chor1 protein lysates prepared in biological triplicate as shown in (g) and (h). Differential protein expression changes were calculated relative to the respective negative control (E3_5 for T-DARPins and sgNTC for sgTBXT). **(j)** Fast preranked GSEA of proteome data with the ARCHS$^4$ Tissue gene collection. Please see panel (d) for a description. **(k)** Fast preranked GSEA of proteome data with the GO_BP gene collection focused on downregulated pathways. Please see panel (e) for a description. **(l)** Fast preranked GSEA of proteome data with the GO_BP gene collection focused on upregulated pathways. Please see panel (e) for a description. **(m)** Circus plot with DEGs categorized according to their target development levels (TDL). The heatmap shows the log2(fold change) of DEGs detected by RNA-seq. DEGs with Tclin level are written out. DEGs also detected by DDA-MS are not shown here but are included in Fig. 4e.

**Supplementary Figure 5**

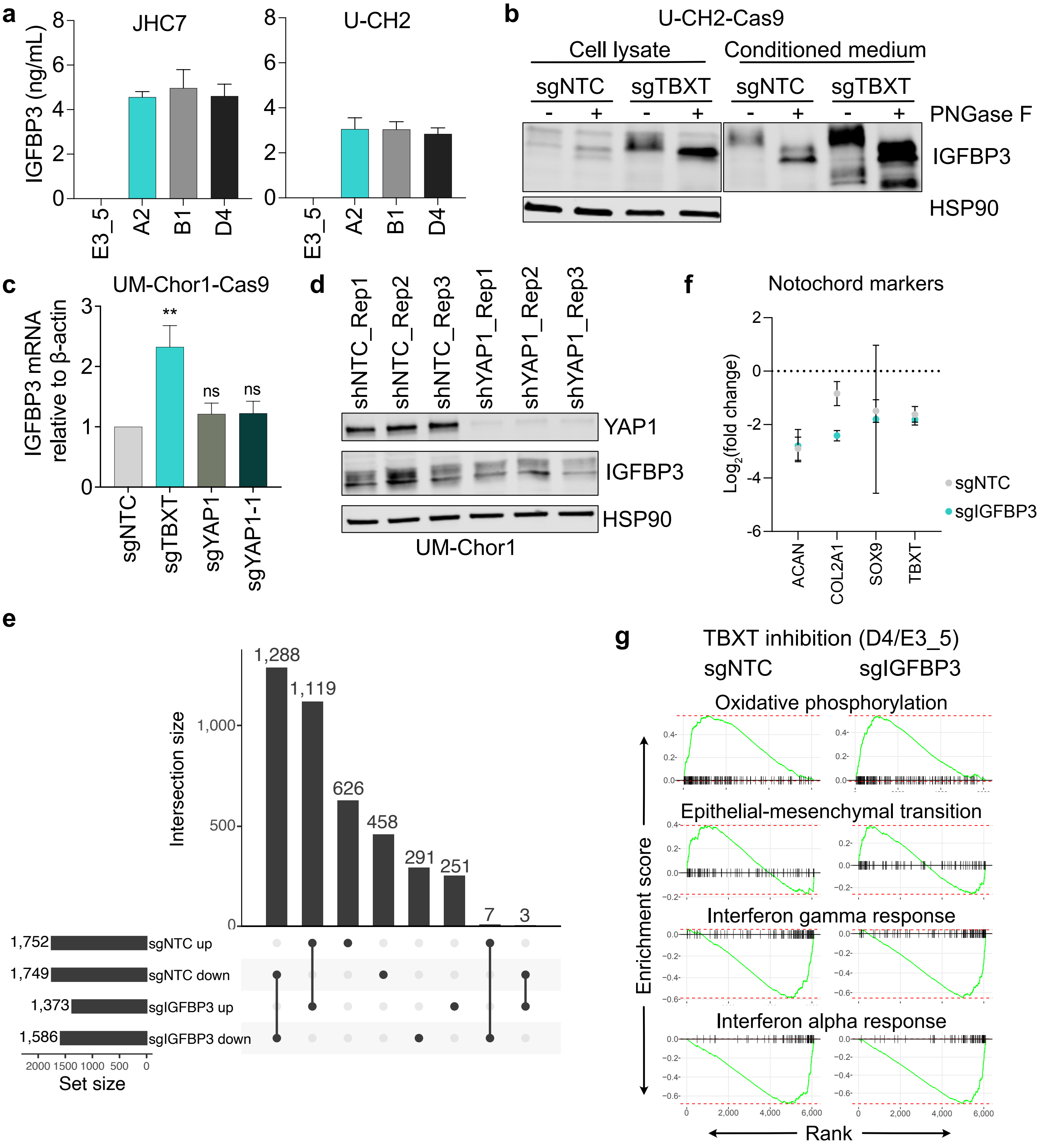


**Supplementary Figure 5. Relationship between IGFBP3 expression and TBXT inhibition. (a)** Cellular protein levels of IGFBP3 quantified by ELISA in JHC7 and U-CH2 cells transduced with DARPins E3_5, A2, B1, or D4. Mean + SEM with two (JHC7 A2) or three (all other conditions) biological replicates. **(b)** Western blot with cell lysates and culture medium of U-CH2-Cas9 cells transduced with sgNTC or sgTBXT. Samples were treated with or without the glycosidase PNGase F. **(c)** *IGFBP3* mRNA levels measured by quantitative RT-PCR in UM-Chor1-Cas9 cells transduced with sgNTC, sgTBXT, or two different sgRNAs targeting YAP1. One-way ANOVA with Dunnett's test for multiple comparisons; mean + SEM of three biological replicates. $^{**}p \leq 0.01$, ns, not significant. **(d)** Western blot of UM-Chor1 cells transduced with an shRNA targeting YAP1 or an shNTC in biological triplicate (Rep1-3). **(e)** UpSet plot showing set and intersection sizes for proteins that are up- or downregulated upon TBXT inhibition with (sgIGFBP3 ) and without (sgNTC) IGFBP3 knock-out. **(f)** Effect of TBXT inhibition (D4/E3_5) on UM-Chor1-Cas9 cells stably expressing sgNTC or sgIGFBP3. Protein levels from DIA-MS graphed as log2(fold change) with upper and lower confidence intervals for four notochord marker genes. **(g)** Fast preranked GSEA plots for the four Hallmark gene sets significantly associated with IGFBP3-dependent response to TBXT inhibition. Green curves show the Kolmogorov-Smirnov random walk, whose maximum deviation from the baseline is depicted with a red dashed line. Positions of genes in each gene set among the ranked proteins are shown on the x-axis with short vertical lines.

**Supplementary Figure 6**

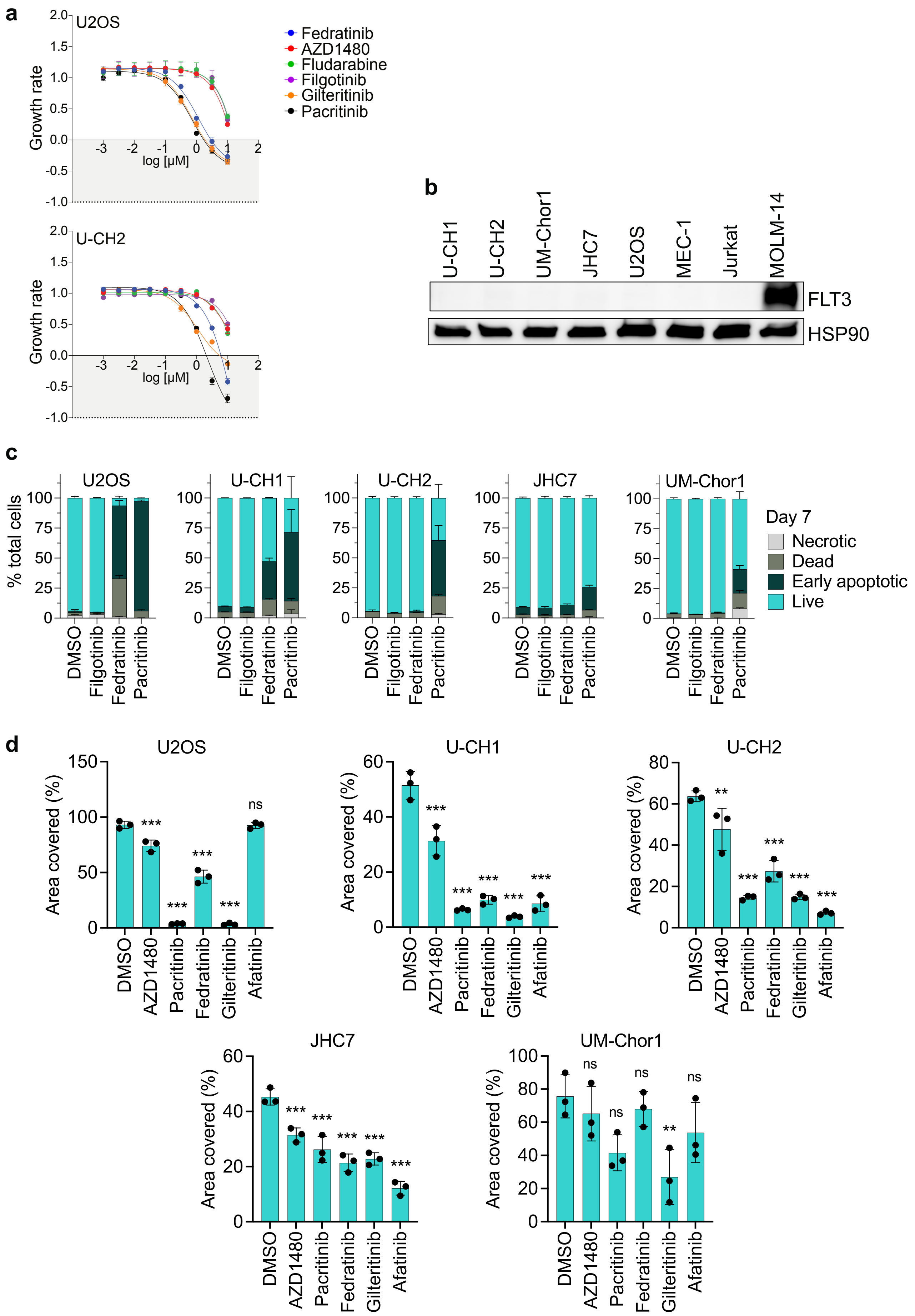

**Supplementary Figure 6. Drug response of chordoma cell lines. (a)** GR analysis of the indicated cell lines after seven days of treatment with the pan-JAK inhibitor filgotinib, the JAK2 inhibitors AZD1480, fedratinib, and pacritinib, the STAT1 inhibitor fludarabine, and the FLT3 inhibitor gilteritinib at concentrations of 10 μM to 1 nM. GR values between 1 and 0 indicate proliferation inhibition, 0 indicates a complete cytostatic effect, and values between 0 and −1 indicate additional cytotoxicity. One of three representative experiments is shown, with each data point representing a technical triplicate. **(b)** Western blot for FLT3 in chordoma cells and the FLT3-positive leukemia cell line MOLM-14**. (c)** Apoptosis measured by flow cytometry after staining with annexin V and 7-AAD of cell lines treated with 3 μM of the indicated drugs for seven days. Mean + SEM of two biological experiments. **(d)** Quantification of colony formation assay shown in Fig. 6f with U2OS and the indicated chordoma cell lines treated for 14 days with 1 μM of the indicated drugs. Mean ± SEM of three independent experiments. One-way ANOVA with Dunnett's test for multiple comparisons; $^{*}p \leq 0.05$, $^{**}p \leq 0.01$, $^{***}p \leq 0.001$, ns, not significant.

### SUPPLEMENTARY METHODS

#### Generation and verification of T-DARPins

Purification of full-length TBXT and the TBXT DNA-binding domain: The purification of the biotinylated recombinant TBXT DBD for ribosome display selection and full-length TBXT for verification after selection was performed by the European Molecular Biology Laboratory Protein Expression and Purification Core Facility, Heidelberg, Germany. Briefly, *E. coli* BL21(DE3) cells were transformed with the pET20b-A(H6)-AviTag plasmid encoding codon-optimized His6-tagged full-length TBXT or the TBXT DBD (amino acids 41–224; sequences provided below). 10 mL overnight culture were added to 1 liter LB broth supplemented with 100 µg/mL carbenicillin. The cultures were grown at 37°C until the optical density at 600 nm reached approximately 0.6. The cultures were then cooled down to 18°C for 30 minutes, expression of TBXT was induced with 0.2 mM IPTG, and cultures were grown overnight at 18°C. Cells were harvested by centrifugation, and the pellets were stored at −20°C. After thawing, the pellets were resuspended in running buffer (50 mM Tris pH 8.0, 250 mM NaCl, 20 mM imidazole) supplemented with 5 mM $MgCl_2$, 10 µg/mL DNAse, 0.5 mg/mL lysozyme, and cOmplete EDTA-free protease inhibitors (Roche). The cells were lysed by two passages through a Microfluidizer. After centrifugation, the supernatant was loaded onto a 5 mL Protino Ni-NTA column (Macherey-Nagel) at 4°C. The Ni-NTA column was washed with running buffer until the A280 nm signal returned to baseline, and the column was eluted with running buffer containing 500 mM imidazole. The elution fractions with full-length TBXT or the TBXT DBD were pooled and dialyzed overnight at 4°C to 50 mM Tris pH 7.0, 50 mM NaCl. The next day, the dialyzed sample was loaded onto a 5 mL HiTrap SP column (Cytiva) pre-equilibrated with 50 mM Tris pH 7.0 and 50 mM NaCl. After washing, the HiTrap SP column was eluted in a gradient to 50 mM Tris pH 7.5 and 1 M NaCl over 12 column volumes. The elution fractions were pooled, concentrated, and injected onto a HiLoad 16/600 Superdex 75 pg column (Cytiva) pre-equilibrated with 20 mM Tris pH 7.5, 300 mM NaCl, 10% glycerol, and 0.1 mM TCEP. The purest protein fractions were pooled, aliquoted, flash-frozen in liquid nitrogen, and stored at −80°C. The identity of the purified protein samples was confirmed by mass spectrometry analysis.

HTRF: Binding of Flag-tagged DARPins to the streptavidin-immobilized, biotinylated TBXT DBD or full-length TBXT was measured using FRET (donor: Streptavidin-Tb cryptate [610SATLB, Cisbio], acceptor: mAb anti-FLAG M2-d2 [61FG2DLB, Cisbio]). Further, HTRF measurements against ‘No Target’ allowed for discrimination of TBXT-specific hits. Experiments were performed at room temperature in white 384-well Optiplate plates (PerkinElmer) using the Taglite assay buffer (Cisbio) at a final volume of 20 µL per well. FRET signals were recorded after 30 minutes using a Varioskan LUX Multimode Microplate (Thermo Scientific). HTRF signals were obtained by dividing the acceptor signal (665 nm) by the donor signal (620 nm) to derive the 665/620 ratio. The background signal was determined using reagents in the absence of DARPins.

ELISA: ELISAs were performed using streptavidin-coated 384-well plates and used for immobilization of the biotinylated TBXT DBD or full-length TBXT at a concentration of 50 nM. Detection of DARPins (1:1,000 dilution of crude extracts) was performed by using a mouse-anti-FLAG M2 monoclonal antibody (dilution 1:5,000; Sigma, F1804) as primary antibody and a goat-anti-mouse antibody conjugated to an alkaline phosphatase (dilution 1:10,000; Sigma, A3562) as secondary antibody. After addition of para-nitrophenyl phosphate, absorbance at 405 nm was determined after 30 minutes. Signals at A540 nm were subtracted as background correction.

Codon-optimized full-length TBXT (5' to 3'):

ATGGCA**CATCACCATCACCATCAC**GGAAGCGGC*GGCCTGAACGATATTTTTGAAGCGCAGAAAATTGAAT*
*GGCATGAA*GGATCCATGAGCAGTCCGGGTACAGAAAGCGCAGGTAAAAGCCTGCAGTATCGTGTTGATCA
TCTGCTGAGCGCAGTTGAAAATGAACTGCAGGCAGGTAGCGAAAAAGGTGATCCGACCGAACGTGAACTG
CGTGTTGGTCTGGAAGAAAGCGAACTGTGGCTGCGTTTTAAAGAACTGACCAATGAAATGATCGTGACCA
AAAATGGTCGTCGCATGTTTCCGGTTCTGAAAGTTAATGTTAGCGGTCTGGACCCGAATGCAATGTATAG
CTTTCTGCTGGATTTTGTGGCAGCAGATAATCACCGTTGGAAATATGTTAATGGTGAATGGGTTCCTGGT
GGTAAACCGGAACCGCAGGCACCGAGCTGTGTTTATATTCATCCGGATAGCCCGAATTTTGGTGCACATT
GGATGAAAGCACCGGTTAGCTTTAGCAAAGTGAAACTGACGAATAAACTGAATGGTGGTGGTCAGATTAT
GCTGAATAGCCTGCATAAATATGAACCGCGTATTCATATTGTTCGTGTTGGT**GGT**CCGCAGCGTATGATT
ACCAGCCATTGTTTTCCAGAAACACAGTTTATTGCAGTTACCGCCTATCAGAACGAAGAAATTACCGCAC
TGAAAATCAAATATAACCCGTTTGCAAAAGCCTTCCTGGATGCAAAAGAACGTAGCGATCATAAAGAAAT
GATGGAAGAACCGGGTGATAGCCAGCAGCCTGGTTATAGCCAGTGGGGTTGGCTGCTGCCTGGTACAAGC
ACCCTGTGTCCGCCTGCAAATCCGCATCCGCAGTTTGGTGGTGCACTGAGCCTGCCGAGCACACATAGCT
GTGATCGTTATCCGACACTGCGTAGCCATCGTAGCAGCCCGTATCCGAGTCCGTATGCACATCGTAATAA
TAGCCCGACCTATAGCGATAATAGTCCGGCATGTCTGAGCATGCTGCAGTCACATGATAATTGGAGTAGC
CTGGGTATGCCTGCACATCCGAGTATGCTGCCGGTTAGCCATAATGCAAGCCCTCCGACCAGCAGCAGTC
AGTATCCGAGCCTGTGGTCAGTTAGCAATGGTGCAGTTACACCGGGTAGCCAGGCAGCCGCAGTTTCAAA
TGGTCTGGGTGCACAGTTTTTTCGTGGTTCACCGGCACATTACACACCGCTGACACATCCGGTTAGCGCA
CCGAGCAGCAGCGGTAGTCCGCTGTATGAAGGTGCAGCAGCAGCAACCGATATTGTGGATAGCCAGTATG
ATGCAGCAGCCCAGGGTCGTCTGATTGCAAGCTGGACACCGGTTTCACCGCCTAGCATGTAATAA

Key for the marked sequence parts:

***6x-His tag***; *AviTag*; codon-optimized TBXT sequence; ***wildtype G177 codon***

Codon-optimized TBXT DBD (5’ to 3’):

ATGGCA***CATCACCATCACCATCAC***GGAAGCGGC*GGCCTGAACGATATTTTTGAAGCGCAGAAAATTGAAT*
*GGCATGAA*GGATCCGAACTGCGTGTTGGTCTGGAAGAAAGCGAACTGTGGCTGCGTTTTAAAGAACTGAC
CAATGAAATGATCGTGACCAAAAATGGTCGTCGCATGTTTCCGGTTCTGAAAGTTAATGTTAGCGGTCTG
GACCCGAATGCAATGTATAGCTTTCTGCTGGATTTTGTGGCAGCAGATAATCACCGTTGGAAATATGTTA
ATGGTGAATGGGTTCCTGGTGGTAAACCGGAACCGCAGGCACCGAGCTGTGTTTATATTCATCCGGATAG
CCCGAATTTTGGTGCACATTGGATGAAAGCACCGGTTAGCTTTAGCAAAGTGAAACTGACGAATAAACTG
AATGGTGGTGGTCAGATTATGCTGAATAGCCTGCATAAATATGAACCGCGTATTCATATTGTTCGTGTTG
GT***GGT***CCGCAGCGTATGATTACCAGCCATTGTTTTCCAGAAACACAGTTTATTGCAGTTACCGCCTATCA
GAACGAAGAAATTACCGCACTGAAAATCAAATATAACTAA

Key for the marked sequence parts:

***6x-His tag***; *AviTag*; codon optimized DBD sequence; ***wildtype G177 codon***

**Cell culture**

UM-Chor1 (kindly provided by the Chordoma Foundation) was maintained in IMDM:RPMI-1640 (4:1, Gibco) with 10% FBS (Sigma-Aldrich), 1% penicillin/streptomycin (P/S, Gibco), and 1x non-essential amino acids (NEAA, Sigma-Aldrich). U-CH1, U-CH2, and U-CH12 (kindly provided by Silke Brüderlein and Peter Möller) were maintained in IMDM:RPMI-1640 (4:1) with 10% FBS, 1% P/S, and 1% L-glutamine (Gibco) on collagen I-coated tissue culture flasks (Corning), and JHC7 (kindly provided by the Chordoma Foundation) in DMEM-F12 (Gibco) with 10% FBS and 1% P/S on collagen I-coated tissue culture flasks. U2OS (kindly provided by Karsten Rippe) and HT-1080 (kindly provided by Marcel Trautmann) were maintained in DMEM (Gibco) with 10% FBS and 1% P/S. MEC-1 (kindly provided by Sascha Dietrich) and Jurkat (kindly provided by Thomas Mercher) were maintained in RPMI with 10% FBS and 1% P/S. HCT-116 (kindly provided by Levi Garraway) and MES-SA [1] were cultured in McCoy's 5a, 10% FBS, and P/S. For experiments, all chordoma cell lines were cultured on collagen I-coated flasks. For UM-Chor1-iDARPin cells, FBS was replaced with tet-free FBS (Takara), and DARPins were induced by the addition of 0.5 µg/mL doxycycline (Th. Geyer) or sterile $H_2O$ as negative control. HEK293T cells were cultured in DMEM with 10% FBS and 1% P/S. All cell lines were cultured under standard conditions (37°C, 5% $CO_2$) and routinely tested for the absence of mycoplasma. Cell line identity was verified using the

Multiplex Cell Authentication Test (Multiplexion) or the Human Cell Line Authentication Service (EurofinsGenomics Germany).

**Vectors**

DARPins (containing an N-terminal MRGS$(H)_6$-tag and a C-terminal Flag-tag) were cloned from the pQIq backbone into the pDONR221 entry vector (Invitrogen) and transferred to the pLenti6.2-V5/DEST lentiviral expression vector (with blasticidin or hygromycin resistance, Invitrogen) and the pLenti CMVtight Puro DEST inducible vector using Gateway Technology (Invitrogen). pLenti CMV rtTA3 Blast (w756-1), pLenti CMV Hygro DEST (w117-1), and pLenti CMVtight Puro DEST were a gift from Eric Campeau (Addgene plasmids #26429, #17454, and #26430). For transient expression in HEK293T cells, DARPins were cloned into the pShuttle CMV-IRES-GFP vector [2] using *SnaBI* and *NotI* restriction enzymes (NEB).

For dual-luciferase reporter assays, the pGL4.34[*luc2P*/SRF-RE/Hygro] vector (Promega, kindly provided by Thordur Oskarsson) encoding the *Firefly* luciferase gene under a minimal promoter was used. To create a TBXT-responsive reporter vector, the serum response factor element was replaced by a DNA sequence containing two palindromic TBXT response elements (2X-T-Resp) [3] using the *Acc65I* and *HindIII*-HF restriction enzymes (NEB) to generate the pGL4.34-2X-T-Resp vector. The 2X-T-Resp forward and reverse oligonucleotides with *Acc65I* and *HindIII* sticky overhangs were synthesized at Sigma-Aldrich with the following sequences (TBXT binding sites underlined): 5'-AGC TAA TTT CAC ACC TAG GTG TGA AAT TCC CGG GAA TTT CAC ACC TAG GTG TGA AAT T-3' (forward) and 5'- GTA CAA TTT CAC ACC TAG GTG TGA AAT TCC CGG GAA TTT CAC ACC TAG GTG TGA AAT T-3' (reverse). The pGL4.73[*hRluc*/SV40] control vector for expression of the *Renilla* luciferase gene was obtained from Promega.

For immunoprecipitation of TBXT variants, the codon-optimized full-length TBXT cDNA with or without a C-terminal HA-tag was synthesized (IDT) and cloned into pLenti6.2-V5/DEST using Gateway Technology. The TBXT-HA cDNAs containing the variants R16L, H171R, G177D, and ΔDBD (removal of amino acids 42–219) were generated in pDONR221 using the QuikChange II Site-Directed Mutagenesis Kit (Agilent). The primer sequences are provided in **Supplementary Table 1**. For transient expression in HEK293T cells, the cDNAs were transferred into the pLEX307 vector (gift from David Root, Addgene plasmid #41392) using Gateway Technology.

For CRISPR/Cas9-mediated knockout, lentiCas9-Blast for Cas9 expression (gift from Feng Zhang, Addgene plasmid #52962) and lentiGuide-Puro (gift from Feng Zhang, Addgene plasmid #52963) and lentiGuide-Hygro (gift from Caroline Goujon, Addgene plasmid #139462) for sgRNA expression were used. The sgRNA sequences (**Supplementary Table 2**) were cloned into the vectors with the *BbsI* restriction enzyme (NEB).

For miR-E-based gene knockdown, we used the SGEP vector (gift from Johannes Zuber, Addgene plasmid #111170). The unmodified SGEP vector contains an shRNA targeting *Renilla* luciferase and was used as a negative control (shLuc). The shRNAs sequences (**Supplementary Table 3**) were cloned into SGEP as PCR amplified 97-mer oligos [4] using *XhoI* and *EcoRI* restriction enzymes (NEB). For shRNA based gene knockdown, we used the pRSI12 vector as described [5] with sequences provided in **Supplementary Table 4**.

**Supplementary Table 1:** Primers used for site-directed mutagenesis

| Name | Primer sequence (5‘ to 3‘) |
|---|---|
| R16L forward | CTCAGCAGATGATCAACAAGATACTGCAGGCTTTTAC |
| R16L reverse | GTAAAAGCCTGCAGTATCTTGTTGATCATCTGCTGAG |
| H171R forward | GACCACCAACACGAACAATACGAATACGCGGTTCATATTTA |
| H171R reverse | TAAATATGAACCGCGTATTCGTATTGTTCGTGTTGGTGGTC |
| G177D forward | CATATTGTTCGTGTTGGTGATCCGCAGCGTATGATTACC |
| G177D reverse | GGTAATCATACGCTGCGGATCACCAACACGAACAATATG |
| ΔDBD forward | GTGATCCGACCGAACGTGAAGCAAAAGAACGTAGC |
| ΔDBD reverse | GCTACGTTCTTTTGCTTCACGTTCGGTCGGATCAC |

**Supplementary Table 2:** sgRNA oligonucleotides

| Name | sgRNA sequence (5‘ to 3‘) |
|---|---|
| sgNTC | AAAAAGCTTCCGCCTGATGG |
| sgTBXT | TGGCTGGTGATCATGCGCTG |
| sgYAP1 | GTGCACGATCTGATGCCCGG |
| sgYAP1-1 | TGCCCCAGACCGTGCCCATG |

| | |
|---|---|
| sgIGFBP3 | CACCAGCTCCGCGCACACGG |

**Supplementary Table 3:** miR-E shRNA oligonucleotides (sense in bold)

| Name | shRNA sequence (5' to 3') |
|---|---|
| shLuc | TGCTGTTGACAGTGAGCG**CAGGAATTATAATGCTTATCT**ATAGTGAAGCCACAGATGTATAGATAAGCATTATAATTCCTATGCCTACTGCCTCGGA |
| shTBXT | TGCTGTTGACAGTGAGCGC**AAGTACAATCCATTTGCAAA**ATAGTGAAGCCACAGATGTATTTTGCAAATGGATTGTACTTATGCCTACTGCCTCGGA |

**Supplementary Table 4:** shRNA oligonucleotides (sense)

| Name | shRNA sequence (5' to 3') |
|---|---|
| shNTC | CAACAAGATGAAGAGCACCAA |
| shYAP | CCCAGTTAAATGTTCACCAAT |

**Lentivirus production and transduction of cells**

Lentivirus was produced by transfection of HEK293T cells with envelope (pMD2.G), packaging (psPAX2), and expression plasmids. pMD2.G and psPAX2 were a gift from Didier Trono (Addgene plasmids #12259 and #12260). The plasmid mix was combined with Opti-MEM (Gibco) and TransIT-LT1 (Mirusbio) transfection reagent and incubated overnight in a 10 cm dish of 70% confluent HEK293T cells. Transfection media was replaced with 3 to 4.5 mL DMEM with 30% FBS and harvested after 24 and 48 hours. Virus-containing medium was passed through a 0.45 µm polyethersulfone membrane filter (PES, Pall) and stored at −80°C or concentrated using PEG-8000 (40 % W/V, VWR International) with 1.2 M NaCl and frozen at −80°C in 1x PBS (Gibco).

For transduction, attached cells in six-well plates were spin-infected for two hours (30°C, 2,000 rpm) in culture medium containing 300 µL unconcentrated virus or 10 µL concentrated virus per well and 4 µg/mL polybrene (Merck Millipore). After overnight incubation, virus-containing medium was removed, cells were trypsinized and washed, plated into cell culture flasks, and, depending on the vector used, subjected to selection with predetermined cell line-specific concentrations of blasticidin, puromycin, or hygromycin for at least four days before the start of experiments. UM-Chor1, JHC7, U-CH1, and U-CH2 cells were transduced with pLenti6.2-V5/DEST encoding E3_5 or T-DARPins and selected with blasticidin. The HEK293T+TBXT cell line was generated with pLenti6.2-V5/DEST encoding TBXT and selected with blasticidin. The UM-Chor1-iDARPin cell lines were generated by transducing cells with pLenti CMV rtTA3 Blast (w756-1), and after selection with blasticidin, the cells were transduced with pLenti CMVtight Puro DEST (w768-1) encoding E3_5 or the T-DARPins and selected with puromycin and blasticidin. For knockout experiments, UM-Chor1-Cas9 and U-CH2-Cas9 cells were generated with lentiCas9-Blast virus and selected with blasticidin. These Cas9-expressing cells were transduced with lentiGuide-Puro-sgRNA virus and selected with puromycin and blasticidin to express single sgRNAs. For combined IGFBP3 knockout and TBXT inhibition, UM-Chor1-Cas9 cells stably expressing sgNTC or sgIGFBP3 were transduced with pLenti CMV Hygro DEST (w117-1) encoding E3_5 or T-DARPin D4 and selected with hygromycin, blasticidin, and puromycin. For IGFBP3 and TBXT double-knockout, UM-Chor1-Cas9 stably expressing sgNTC and sgIGFBP3 were transduced with lentiGuide-Hygro containing sgNTC and sgTBXT and selected with hygromycin, blasticidin, and puromycin. For knockdown experiments, UM-Chor1 or U-CH2 cells were transduced with either pRSI12 containing shNTC or shYAP or SGEP containing shLuc or shTBXT and selected with puromycin.

**Western blotting**

**Supplementary Table 5:** Primary and secondary antibodies used for western blotting

| Name | Source species | Dilution | Company | Article # |
|---|---|---|---|---|
| Anti-TBXT (D2Z3J) | Rabbit | 1:1,000 in 5% BSA/TBST | Cell Signaling | 81694 |
| Anti-FLAG | Rabbit | 1:400 in 5% BSA/TBST | Sigma-Aldrich | F7425 |
| Anti-β-actin (AC-15) | Mouse | 1:2,000 in 5% BSA/TBST | Sigma-Aldrich | A1978 |
| Anti-β-actin | Rabbit | 1:1,000 in 5% BSA/TBST | Cell Signaling | 4967 |
| Anti-HSP90 (F-8) | Mouse | 1:5,000 in 5% BSA/TBST | Santa Cruz | sc-13119 |
| Anti-YAP (D8H1X) | Rabbit | 1:1,000 in 5% BSA/TBST | Cell Signaling | 14074 |
| Anti-IGFBP3 (EPR18680-153) | Rabbit | 1:1,000 in 5% BSA /TBST | ABCAM | ab193910 |
| Anti-IGFBP3 (D1U9C) | Rabbit | 1:1,000 in 5% BSA/TBST | Cell Signaling | 25864 |
| Anti-ISG15 (EPR3446) | Rabbit | 1:1,000 in 5% BSA/TBST | Abcam | ab133346 |

| | | | | |
|---|---|---|---|---|
| Anti-JAK1 | Rabbit | 1:1,000 in 5% BSA/TBST | Cell Signaling | 3332 |
| Anti-JAK2 (D2E12) | Rabbit | 1:1,000 in 5% BSA/TBST | Cell Signaling | 3230 |
| Anti-JAK3 | Rabbit | 1:1,000 in 5% BSA/TBST | Cell Signaling | 3775 |
| Anti-TYK2 (D4I5T) | Rabbit | 1:1,000 in 5% BSA/TBST | Cell Signaling | 14193 |
| Anti-STAT1 (D1K9Y) | Rabbit | 1:1,000 in 5% BSA/TBST | Cell Signaling | 14994 |
| Anti-STAT2 (D9J7L) | Rabbit | 1:1,000 in 5% BSA/TBST | Cell Signaling | 72604 |
| Anti-STAT3 (124H6) | Mouse | 1:1,000 in 5% BSA/TBST | Cell Signaling | 9139 |
| Anti-STAT4 (C46B10) | Rabbit | 1:1,000 in 5% BSA/TBST | Cell Signaling | 2653 |
| Anti-STAT5 (D206Y) | Rabbit | 1:1,000 in 5% BSA/TBST | Cell Signaling | 94205 |
| Anti-STAT6 (D3H4) | Rabbit | 1:1,000 in 5% BSA/TBST | Cell Signaling | 5397 |
| Anti-IRF9 (D2T8M) | Rabbit | 1:1,000 in 5% BSA/TBST | Cell Signaling | 76684 |
| Anti-EGFR | Rabbit | 1:1,000 in 5% BSA/TBST | Cell Signaling | 4267 |
| Anti-pEGFR(Y1068) | Rabbit | 1:1,000 in 5% BSA/TBST | Cell Signaling | 3777 |
| Anti-mouse IgG (H+L) DyLight 680 Conjugate | Goat | 1:15,000 in 5% milk/TBST | Cell Signaling | 5470 |
| Anti-rabbit IgG (H+L) DyLight 800 4X PEG Conjugate | Goat | 1:15,000 in 5% milk/TBST | Cell Signaling | 5151 |
| Anti-rabbit IgG, HRP-linked | Goat | 1:100,000 in 5% milk/TBST | Cell Signaling | 7074 |
| Anti-mouse IgG, HRP-linked | Horse | 1:100,000 in 5% milk/TBST | Cell Signaling | 7076 |

**Mass spectrometry sample preparation, data acquisition, and preprocessing**

AP-MS: The eluate (35 µl) of each IP sample was run for 0.5 cm into an SDS-PAGE and the entire piece was cut out and digested with trypsin according to Shevchenko et al. [6] adapted to a DigestPro MSi robotic system (INTAVIS Bioanalytical Instruments AG). The LC-MS/MS analysis was carried out on an Ultimate 3000 UPLC system (Thermo Fisher Scientific) directly connected to a Q-Exactive HF-X Orbitrap mass spectrometer for a total of 90 minutes. Peptides were online desalted on a trapping cartridge (Acclaim PepMap300 C18, 5 µm, 300 Å wide pore; Thermo Fisher Scientific) for three minutes using a 30 µL/minute flow of 0.05% TFA in water. The analytical multistep gradient (300 nL/minute) was performed using a nanoEase MZ Peptide analytical column (300 Å, 1.7 µm, 75 µm x 200 mm, Waters) using solvent A (0.1% formic acid in water) and solvent B (0.1% formic acid in 80% acetonitrile with 20% water). For 50 minutes the concentration of B was linearly ramped from 2% to 25% and to 40% within the next 10 minutes, followed by a quick ramp to 95%, after five minutes the concentration of B was lowered to 2% and a 20 minute equilibration step appended. Eluting peptides were analyzed in the mass spectrometer using data dependent acquisition (DDA) mode. A full scan at 120k resolution (375-1500 m/z, 3e6 AGC target, 54 ms maxIT) was followed by up to 10 MS/MS scans. Peptide features were isolated with a window of 1.6 m/z, fragmented using 27% NCE. Fragment spectra were recorded at 30k resolution (1.5e5 AGC target, 54 ms maxIT). Unassigned and singly charged eluting features were excluded from fragmentation and dynamic exclusion was set to 10 seconds. Each sample was followed by a wash run (60 minutes) to minimize carry-over between samples. Instrument performance throughout the course of the measurement was monitored by regular (approximately one per 48 hours) injections of a standard sample and an in-house shiny application. Data preprocessing was carried out by MaxQuant (2.1.4.0) [7] using an organism specific database extracted from Uniprot.org (human reference database with one protein sequence per gene, containing 20,591 unique entries from March 1$^{st}$, 2023 and nine DARPin sequences). Settings were left as default with the following adaptions: Match between runs (MBR) was enabled to transfer peptide identifications across RAW files based on accurate retention time and m/z. Fractions were set in a way that MBR was only performed within replicates. Label free quantification (LFQ) was enabled with default settings. The iBAQ-value [8] generation was enabled.

DDA-MS: Proteins (10 µg) were run for 0.5 cm into an SDS-PAGE and the entire piece was cut out and digested using trypsin according to Shevchenko et al. [6] adapted to a DigestPro MSi robotic system (INTAVIS Bioanalytical Instruments AG). The LC-MS/MS analysis was carried out on an Ultimate 3000 UPLC system (Thermo Fisher Scientific) directly connected to an Orbitrap Exploris 480 mass spectrometer for a total of 150 minutes. Peptides were online desalted on a trapping cartridge (Acclaim PepMap300 C18, 5 µm, 300 Å wide pore; Thermo Fisher Scientific) for three minutes using 30 µL/minute flow of 0.05% TFA in water. The analytical multistep gradient (300 nL/minute) was performed using a nanoEase MZ Peptide analytical column (300 Å, 1.7 µm,

75 µm x 200 mm, Waters) using solvent A (0.1% formic acid in water) and solvent B (0.1% formic acid in acetonitrile). For 132 minutes the concentration of B was linearly ramped from 4% to 30%, followed by a quick ramp to 80%, after two minutes the concentration of B was lowered to 2% and a 10 minute equilibration step appended. Eluting peptides were analyzed in the mass spectrometer using DDA mode. A full scan at 120k resolution (380-1400 m/z, 300% AGC target, 45 ms maxIT) was followed by up to two seconds of MS/MS scans. Peptide features were isolated with a window of 1.4 m/z, fragmented using 26% NCE. Fragment spectra were recorded at 15k resolution (100% AGC target, 54 ms maxIT). Unassigned and singly charged eluting features were excluded from fragmentation and dynamic exclusion was set to 35 seconds. Each sample was followed by a wash run (40 minutes) to minimize carry-over between samples. Instrument performance throughout the course of the measurement was monitored by regular (approximately one per 48 hours) injections of a standard sample and an in-house shiny application. Data preprocessing was carried out by MaxQuant (2.1.4.0) [7] using an organism specific database extracted from Uniprot.org (human reference database with one protein sequence per gene, containing 79,038 unique entries from January 3$^{rd}$, 2022 and four DARPin sequences). Settings were left at default with the following adaptions: MBR was enabled to transfer peptide identifications across RAW files based on accurate retention time and m/z. Fractions were set in a way that MBR was only performed within replicates. LFQ was enabled with default settings. Separate parameter groups were assigned for DARPin (E3_5, A2, B1, D4) and TBXT knock-out (sgNTC, sgTBXT) samples, including separate LFQ normalization. The iBAQ-value [8] generation was enabled.

DIA-MS: Proteins (10 µg) were digested with trypsin using an AssayMAP Bravo liquid handling system (Agilent technologies) running the autoSP3 protocol according to Müller et al [9]. The LC-MS/MS analysis was carried out on an Ultimate 3000 UPLC system (Thermo Fisher Scientific) directly connected to an Orbitrap Exploris 480 mass spectrometer for a total of 120 minutes. Peptides were online desalted on a trapping cartridge (Acclaim PepMap300 C18, 5 µm, 300 Å wide pore; Thermo Fisher Scientific) for three minutes using 30 µL/minute flow of 0.05% TFA in water. The analytical multistep gradient (300 nL/minute) was performed using a nanoEase MZ Peptide analytical column (300 Å, 1.7 µm, 75 µm x 200 mm, Waters) using solvent A (0.1% formic acid in water) and solvent B (0.1% formic acid in acetonitrile). For 102 minutes the concentration of B was linearly ramped from 4% to 30%, followed by a quick ramp to 78%, after two minutes the concentration of B was lowered to 2% and a 10 minute equilibration step appended. Eluting peptides were analyzed in the mass spectrometer using data independent acquisition (DIA) mode.

A full scan at 120k resolution (380-1400 m/z, 300% AGC target, 45 ms maxIT) was followed by data independent MS2 acquisition covering a mass range of 400-1000 m/z via 47 DIA windows of variable width with one Da overlap and 28% NCE. Fragment spectra were recorded at a resolution of 30k (1000% AGC target, 54 ms maxIT). Each sample was followed by a wash run (40 minutes) to minimize carry-over between samples. Instrument performance throughout the course of the measurement was monitored by regular (approximately one per 48 hours) injections of a standard sample and an in-house shiny application. Preprocessing of DIA RAW files was performed with Spectronaut (Biognosys, version 17.1.221229.55965) in directDIA+ (deep) library-free mode. Default settings were applied with the following adaptions: Within the Pulsar Search in Peptides, the Max Peptide Length was set to 35, in Result Filters, the Peptide Charge was enabled and the Max Charge set to 6 and the Min Charge set to 2. Within DIA Analysis under Identification the Precursor PEP, Cutoff was set to 0.01, the Protein Qvalue Cutoff (Run) set to 0.01, and the Protein PEP Cutoff set to 0.05. In Quantification, the Proteotypicity Filter was set to Only Protein Group Specific, the Quantification window was set to Not Synchronized, and the Major Group Quantity was set to Sum peptide quantity. The data was searched against the human proteome from Uniprot (human reference database with one protein sequence per gene, containing 20,591 unique entries from March 1$^{st}$, 2023), the contaminants FASTA from MaxQuant (246 unique entries from December 22$^{nd}$, 2022) and the sequences for E3_5 and D4.

**Quantitative and regular RT-PCR**

**Supplementary Table 6:** Primer sequences used for quantitative RT-PCR.

| Name | Primer sequence (5‘ to 3‘) |
|---|---|
| SOX9 forward | AGCGAACGCACATCAAGAC |
| SOX9 reverse | CTGTAGGCGATCTGTTGGGG |
| TBXT forward | TATGAGCCTCGAATCCACATAGT |
| TBXT reverse | CCTCGTTCTGATAAGCAGTCAC |
| ACTB forward | CATGTACGTTGCTATCCAGGC |
| ACTB reverse | CTCCTTAATGTCACGCACGAT |

For the amplification of DARPins from xenografted tumors, mRNA was purified from the U-CH1 tumors using a Qiagen RNeasy Mini kit. 500 ng RNA was reverse transcribed using the High-Capacity cDNA Reverse Transcription Kit (Applied Biosystems). The cDNA was diluted into 100 µL of nuclease free water. 10 ng of cDNA was amplified in a 50 µL reaction with 0.5 µL Velocity polymerase (Meridian Bioscience, #BIO-21098), 10 µL Hi-Fi Buffer, 1 µL dNTPs (Qiagen, #201912), 1 µL DMSO, 1 µL forward and reverse primer, and amplified using the following program: 98°C – 3 mins, 98°C – 20s / 68°C – 30s / 72°C – 1min for 30 cycles, 72°C – 5 min. The primer sequences were as follows: DARPin-forward 5'-ATGGGCATGAGAGGATCGC-3' and DARPin-reverse 5'-TGTCGTCGTCATCCTTGTAGT-3'.

**RNA-seq data preprocessing**

Reads were processed using the RNA-seq workflow 1.3.0 developed by the DKFZ Omics IT and Data Management Core Facility (https://github.com/DKFZ-ODCF/RNAseqWorkflow). First, FASTQ reads were aligned to the 1,000 Genomes Project Phase II Reference Genome (hs37d5) based on NCBI GRCh37 via two-pass alignment using STAR 2.5.3a [10]. The STAR index was generated from the 1,000 Genomes assembly and GENCODE Version 19 gene models with a sjdbOverhang of 200. Alignment call parameters were as follows:

| Parameter | Value |
|---|---|
| --alignIntronMax | 1100000 |
| --alignIntronMin | 20 |
| --alignMatesGapMax | 1100000 |
| --alignSJstitchMismatchNMax | 5 -1 5 5 |
| --alignSJDBoverhangMin | 3 |
| --chimJunctionOverhangMin | 15 |
| --chimScoreMin | 1 |
| --chimScoreJunctionNonGTAG | 0 |
| --chimSegmentMin | 15 |
| --chimSegmentReadGapMax | 3 |
| --clip3pAdapterSeq | AGATCGGAAGAGCACACGTCTGAACTCCAGTCA |
| --genomeLoad | NoSharedMemory |
| --limitBAMsortRAM | 100000000000 |
| --outBAMsortingThreadN | 1 |
| --outSAMstrandField | intronMotif |
| --outSAMtype | BAM Unsorted SortedByCoordinate |
| --outSAMunmapped | Within KeepPairs |
| --outFilterMismatchNmax | 5 |
| --outFilterMismatchNoverLmax | 0.3 |
| --outFilterMultimapNmax | 1 |
| --readFilesCommand | gunzip -c |

| | |
|---|---|
| --runThreadN | 8 |
| --sjdbOverhang | 200 |
| --twopass1readsN | -1 |
| --twopassMode | Basic |

Duplicate marking of the resultant main alignment file was done with sambamba 0.6.5 [11]. The chimeric file was sorted using samtools 1.6 [12]. BAM indexes were generated using sambamba 0.6.5. Quality control analysis was performed using samtools flagstat and the rnaseqc tool version 1.1.8 [13]. Depth of coverage analysis for rnaseqc was turned off. Gene-specific read counting was performed using featureCounts version 1.5.1 [14] over exon features based on the GENCODE 19 gene models. Only unique alignments were counted. Strand specific counting was also used. A custom script was used to calculate RPKM and TPM expression values. For total library abundance calculations, all genes on chromosomes X, Y, MT as well as rRNA and tRNA genes were omitted, as they are likely to introduce library size estimation biases.

**Determination of viable cell numbers**

To assess the effect of DARPin expression on chordoma cell number, 30,000 cells per well were seeded in six-well plates, the medium was changed once a week, and cells were detached and counted on day 21 using a Vi-CELL counter (Beckman) and trypan blue exclusion.

To assess the effect of DARPin expression on cell proliferation, 3,000 (HCT-116 and HT-1080), or 1,500 (MES-SA) cells per well were seeded in 96-well plates and cell viability was assessed after different incubation times with the CellTiter 96® AQueous One Solution Cell Proliferation Assay (Promega) according the manufacturer's instructions. Absorbance at 490 nm was determined with an iMark Microplate Reader.

**Cell cycle analysis**

Twelve days after transduction, 0.5 to 1 x $10^6$ UM-Chor-1 cells were trypsinized, washed using 1 mL of cold PBS containing 0.1% BSA, and centrifuged with 200 x g for five minutes at 4°C. Cell pellets were resuspended in 100 µL cold PBS/0.1% BSA and fixed by drop-wise adding 900 µL of ice-cold 80% ethanol. Cells were stored at −20°C for a minimum of six hours, washed with 1 mL cold PBS/0.1% BSA, and centrifuged with 200 x g for 10 minutes at 4°C. After two washing steps, the cells were resuspended in 500 µL PBS containing 50 µg/mL PI (Invitrogen) and 200 µg/mL RNAse A (Qiagen) and were incubated for 30 minutes at 37°C. The analysis was performed by flow cytometry using a FACSCelesta instrument (BD Biosciences), and the acquired data were analyzed using FlowJo (BD Biosciences).

**Apoptosis analysis**

UM-Chor1, JHC7, U-CH2, and U-CH12 cells were transduced with DARPins, selected, and reseeded without selection antibiotic after four days. After six days, i.e., ten days after transduction, adherent and floating cells were harvested, washed in PBS, and stained with annexin V and 7-AAD (FITC Annexin V Apoptosis Detection Kit I, BD Biosciences) according to the manufacturer's instructions. Cells were analyzed by flow cytometry using an LSR Fortessa instrument (BD Biosciences). For the assessment of drug sensitivity, chordoma ($2 \times 10^5$) or U2OS ($1 \times 10^5$) cells were seeded in collagen-coated t25 flasks and 24 hours later were incubated with DMSO or 3 µM drug for 4 or 7 days. Adherent and floating cells were harvested, washed in PBS, and stained with annexin V (Invitrogen, #R37174) and 7-AAD (BD Biosciences, #559925) and analyzed by flow cytometry using a FACSCelesta instrument (BD Biosciences) after four or seven days of treatment. Flow cytometry data were analyzed with FlowJo (version 10.10).

**Immunofluorescence**

UM-Chor1 cells were analyzed 10 days after infection in 12-well collagen-coated plates (Corning). The medium was removed and cells were washed with cold DPBS and fixed with the Image-iT Fixation/Permeabilization Kit according to manufacturer's instructions (Invitrogen, #R37602). Cells were stained with mouse Penta-His Alexa Fluor 488 antibody (1:250 in 1x DPBS/3% BSA, #35310, Qiagen) for one hour at room temperature, washed three times with 1x DPBS, and stained for one hour with DAPI (0.125 µg/mL) and Texas Red-X Phalloidin (1:400, #T7471, Invitrogen) in 1x DPBS/3% BSA. Cells were washed three times with 1x DPBS and then imaged using a Lionheart FX automated microscope (BioTek). Spheroids were processed for confocal microscopy using a protocol modified from [xx]. Briefly, spheroids were centifuged at 50 g for 3 minutes, and after removing the medium, were transferred to 1% BSA pre-coated 2 mL round bottom tubes. They were washed two times with cold 1x DPBS/1% BSA, fixed at 4°C using 1x DPBS/4% PFA for 45 minutes, permeabilized at 4° using cold 1x DPBS/0.1% Tween-20 for 10 minutes, blocked at 4°C using 1x DPBS/0.1% Triton/0.2% BSA for 15 minutes, and then transferred to a 24-well plate (precoated with 1% BSA). The spheroids were incubated with the primary antibodies mouse anti-Ki-67 (1:200 in 1x DPBS/0.1% Triton/0.2% BSA, #9449, Cell Signaling Technology) or rabbit anti-TBXT (1:200 in 1x DPBS/0.1% Triton/0.2% BSA, #81694, Cell Signaling Technology) overnight at 4°C with gentle rocking. They were then washed 3 times with 1x DPBS/0.1% Triton/0.2% BSA for 2 hours, stained with the secondary antibodies goat anti-rabbit IgG 488 (1:1,000 in 1x DPBS/0.1% Triton/0.2% BSA, #A32731, Invitrogen) or goat anti-mouse IgG 488 (1:1,000 in 1x DPBS/0.1% Triton/0.2% BSA, #A32723, Invitrogen) overnight at 4°C with gentle rocking, were then washed 3 times with 1x DPBS/0.1% Triton/0.2% BSA for 2 hours, incubated with DAPI (0.125 µg/mL) and Texas Red-X Phalloidin (1:400) in 1x DPBS/0.1% Triton/0.2% BSA for 30 minutes, washed 3 times in 1x DPBS/0.1% Triton/0.2% BSA (10 minutes), and then incubated in clearing solution (60% glucose, 2M fructose) for 20 minutes prior to acquisition. Spheroids were imaged using an A1R MP confocal microscope (Nikon).

**IGFBP3 ELISA**

10 µg chordoma cell lysate and a serial dilution of recombinant IGFBP3 protein (0.1 µg/mL to 74.97 pg/mL, Millipore) to create a standard curve for quantification was incubated overnight in an ELISA plate coated with anti-human IGFBP3 antibody (Millipore, RAB0235) at 4°C. The ELISA and quantification were performed according to the manufacturer's instructions. Absorbance was recorded at 450 nm using a Victor X3 plate reader (Perkin Elmer).

**IGFBP3 glycosylation and secretion analysis**

UM-Chor1-Cas9 and U-CH2-Cas9 cells were transduced with sgNTC or sgTBXT lentivirus, selected for 14 days, and serum starved for 72 hours. Protein lysate was prepared as described above. Conditioned medium (CM; 11 mL per t75 flask) was collected, centrifuged with 300 g at 4°C for five minutes to remove cell debris, and stored at −80°C. To isolate secreted proteins, we used an adapted protocol employing trichloroacetic acid (TCA, SERVA Electrophoresis) and sodium deoxycholate (DOC, #D6750, Sigma-Aldrich) [15]. CM was thawed on ice, mixed with DOC to 1% (v/v), incubated on ice for 30 minutes, mixed with TCA to 7.5% (v/v), and incubated on ice for 60 minutes. The processed CM was transferred to 1.5 mL tubes and centrifuged with 15,000 g at 4°C for 20 minutes. The supernatant was discarded, and protein pellets were washed with 1 mL ice-cold acetone, gently vortexed, and placed at −20°C for five minutes. The tubes were centrifuged with 15,000 g at 4°C for 20 minutes, the supernatant was discarded, and the washing step was repeated. After final supernatant removal, cleaned pellets were air-dried in a chemical hood for 30 minutes and stored at −80°C. For the PNGase F digestion, two pellets per condition were dissolved in a total of 50 µL 1x PNGase F buffer (Thermo Fisher) and incubated for three hours at 4° in a tube shaker (750 rpm, Eppendorf) with mixing by pipette every 30 minutes. Residual TCA was neutralized using 5 µL 1M Tris-HCL (pH 8), and samples were frozen at −80°. PNGase F digestion was performed according to the manufacturer's instructions with a few minor modifications using a PNGase F Glycan Cleavage Kit (Thermo Fisher). For total cell lysate digestion, 25 µg protein per condition were mixed with 4 µL 10x PNGase F buffer and 0.5 µL PNGase F and adjusted to a total volume of 40 µL with HPLC-grade water. The reaction was incubated for one hour at 50°C to digest glycosylated residues. For undigested control samples, PNGase F was omitted. For CM samples, equal volumes of isolated protein were digested to normalize across conditions. Therefore, 10 µL of CM-isolated protein was mixed with 4 µL 10x PNGase F buffer and 0.5 µL PNGase F and adjusted to a total volume of 40 µL with HPLC-grade water. For subsequent SDS-PAGE and western blotting, total cell lysate samples were normalized by mass (10 µg/sample), and CM samples were normalized by volume (33.75 µL of reaction) before loading.
